# Group Heteroscedasticity - A Silent Saboteur of Power and False Discovery in RNA-Seq Differential Expression

**DOI:** 10.1101/2024.04.01.587633

**Authors:** Suvo Chatterjee, Arindam Fadikar, Vrushab Hanumesh, Siddhant Sunil Meshram, Roger S Zoh, Siyuan Ma, Ganesan Arunkumar, Himel Mallick

## Abstract

Despite the availability of several high-profile, state-of-the-art methods, analyzing bulk RNA-Seq data continues to face significant challenges. Evidence from recent studies has highlighted that popular differential expression (DE) tools, such as edgeR and DESeq2, are susceptible to an alarmingly high false discovery rate (FDR). These studies suggest that the FDR inflation observed in these models could be attributed to issues such as violations of parametric assumptions or an inability to effectively handle outliers in the data. Here, we argue that group heteroscedasticity can also contribute to this elevated FDR, a phenomenon largely overlooked by the research community. We introduce a novel statistical model, Robseq, designed for effective per-feature modeling in differential analysis, particularly when the assumption of group homoscedasticity is unmet. Robseq utilizes well-established statistical machinery from the robust statistics literature, including M-estimators to robustly estimate gene expression level changes and Huber-Cameron variance estimators to calculate robust standard errors in heteroscedastic settings. Additionally, it incorporates a degrees of freedom adjustment for the Welch t-statistic, based on Bell-McCaffrey’s recommendation, for inferential purposes, effectively addressing the problem of FDR inflation in RNA-Seq differential expression. Through detailed simulations and comprehensive benchmarking, we show that Robseq successfully maintains the false discovery and type-I error rates at nominal levels while retaining high statistical power compared to well-known DE methods. Analysis of population-level RNA-Seq data further demonstrates that Robseq is capable of identifying biologically significant signals and pathways implicated in complex human diseases that otherwise cannot be revealed by published methods. The implementation of Robseq is publicly available as an R package at https://github.com/schatterjee30/Robseq.

## 1 Introduction

Traditional bulk RNA sequencing (RNA-Seq) provides a revolutionary way to uncover the transcriptomic landscape of health and disease by measuring the gene expression of thousands of genes simultaneously over tens of thousands of cells using ultra-high-throughput sequencing ^68,71^. Naturally, RNA-Seq has emerged as a universal and established technology in modern biology over the last decade or so for unraveling the molecular basis of biological processes ^9,8^, as highlighted by increasingly common translational and clinical studies utilizing transcriptome profiling across a variety of disease areas and therapeutic outcomes ^37,26,60,69,42,22^. Diverse applications of RNA-Seq include patient stratification, disease subtype identification, biomarker discovery, and mechanism of action study, among others ^17,63^.

A key analytic task in RNA-Seq studies is identifying genes with differential expression (DE) patterns, commonly referred to as DE genes, which exhibit changes in expression levels across two or more conditions. In the past decade, numerous statistical approaches have been proposed to perform DE analysis of RNA-Seq data. However, despite the availability of a plethora of high-profile methods proposed for DE analysis, a clear consensus on the most efficient approach has yet to be reached. These methods broadly fall under two categories: parametric and nonparametric. The parametric methods rely on assumptions about probability models either based on raw counts using a discrete probability distribution or on transformed counts using a continuous probability distribution. Among the parametric models, the sub-categories include, (i) Poisson- and generalized Poisson-based model, such as DEGseq ^67^, deGPS ^11^, and TSPM ^3^, (ii) methods based on a negative binomial (NB) model, such as edgeR ^53^, robust edgeR ^76^, quasi-likelihood edgeR ^41^, DESeq2 ^40^, EBSeq ^34^, and NBPSeq ^16^, (iii) methods based on linear models, such as limma-trend and voom ^33^, and (iv) methods based on a beta-binomial model, such as BBSeq ^77^. In contrast, the nonparametric methods do not assume any particular parametric model and include methods such as NOISeq ^59^, SAMseq ^35^, and dearseq ^21^, among others.

Despite the remarkable advancements made, RNA-Seq DE analysis still presents some challenges, perhaps the most important being the control of the false discovery rate (FDR) at the nominal level. The FDR pertains to the expected ratio of false positive classifications (false discoveries) to the total number of significant findings. The absence of robust FDR control leads to an overabundance of false positive results, thereby diminishing the reproducibility of research outcomes. This is a critical gap to address from a public health standpoint, as an excessive number of false positives introduces ambiguity during interpretation and misallocates valuable resources based on false signals. Several recent studies ^55,12,38,21,54,2^ have shed light on the issue. Among these, many emphasized the problem of FDR inflation in state-of-the-art models such as edgeR and DESeq2 ^55,12,38,21,54^ while others examined the severeness of this phenomenon in both small and large sample sizes ^38,21^. Additionally, studies have emerged that reached a similar conclusion when analyzing long non-coding RNA expression data ^2^. This implies that investigators are compelled to choose one of the popular DE tools for their day-to-day RNA-Seq data analysis without much guidance on their effectiveness in controlling the FDR.

The collective evidence in the literature thus demonstrates that widely employed DE methods, such as edgeR and DESeq2, exhibit susceptibility to FDR inflation across realistically complex scenarios. These investigations ^55,12,38,21,54,2^ also provide reasoning that the observed lack of FDR control in these gold standard models might stem from issues such as inaccurate distributional assumptions, like the negative binomial distribution for modeling the sequence counts ^25^, or the inability of the models to account for outliers within the data effectively. Another potential cause of FDR inflation could be the complete disregard for heteroscedasticity by popular DE methods despite its omnipresence in RNA-Seq data (**Fig. 1**). Notably, heteroscedasticity, characterized by varying dispersion among groups, can distort the FDR by challenging the fundamental assumptions of statistical tests, which typically assume homoscedasticity (i.e., equal variability across diverse conditions or groups). This uneven variability compromises standard error calculations in hypothesis testing, leading to biased and misaligned test statistics no longer approximately centered at zero, as expected under the null and homoscedasticity assumption. While heteroscedasticity in the context of RNA-Seq data has not been widely studied, it has been reported in the literature ^15,39,10,50^. For instance, the study by Ran et al. ^50^ identified challenges in limma voom’s capability to model variability accurately under scenarios involving heteroscedasticity. Another study by Chen et al. ^10^ highlighted unequal group variance in single-cell data and emphasized the under-utilization of tests accounting for varying variances. More recently, a study by You et al. ^72^ highlighted that group heteroscedasticity is present in pseudo-bulk data and subsequently proposed a statistical model to account for it. However, the extent to which heteroscedasticity impacts false discovery and power, especially in large samples, remains to be investigated. We propose that accounting for group heteroscedasticity in bulk RNA-Seq data analysis can address the issue of FDR inflation without compromising statistical power. This insight drives our effort in the development of a new statistical pipeline that can effectively conduct DE analysis within a heteroscedastic framework while concurrently upholding statistical power and control over false discoveries across a wide variety of settings.

**Figure 1:**
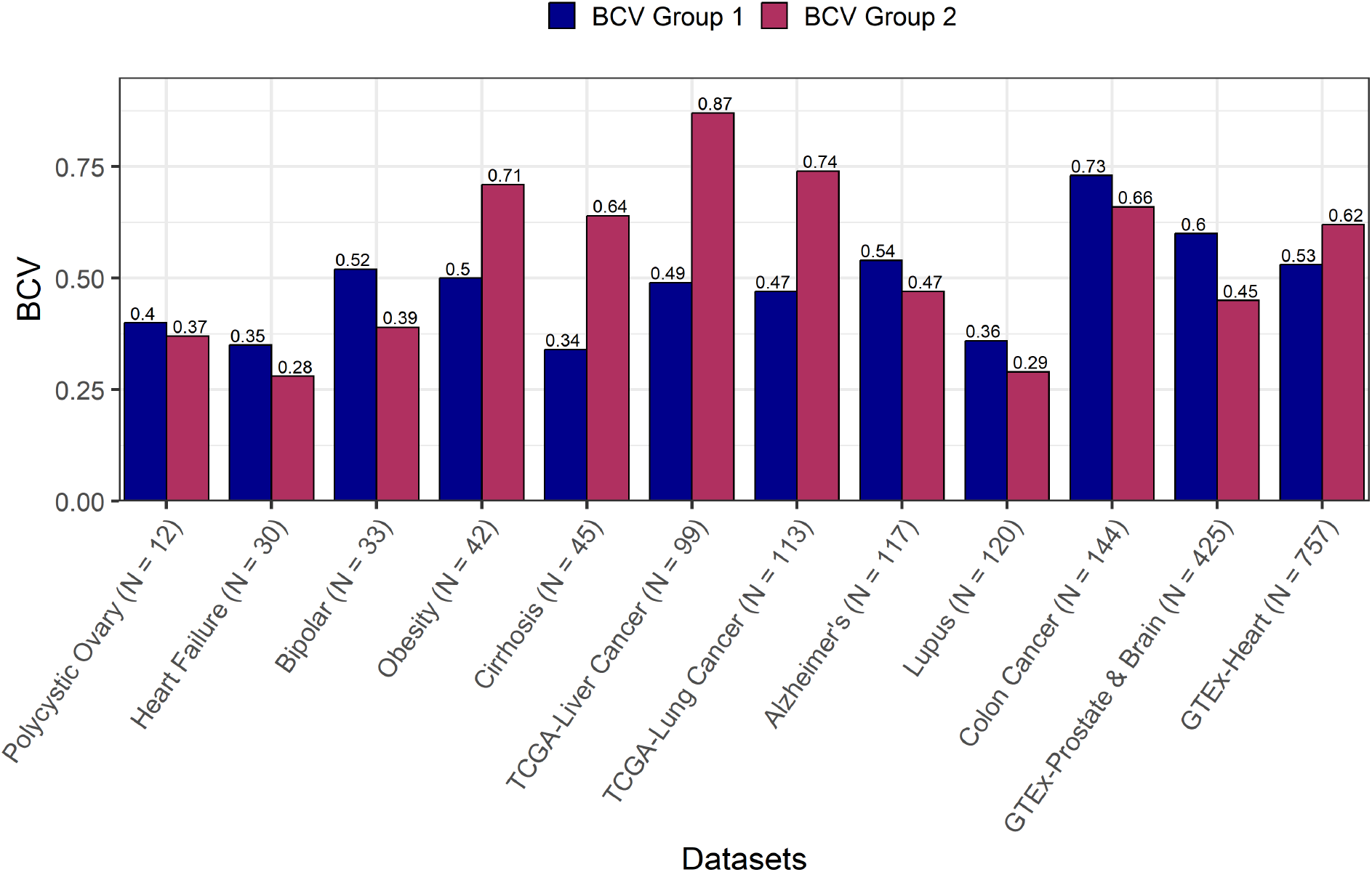
Group heteroscedasticity is commonly observed in bulk RNA-Seq data. We calculated group-specific biological coefficient of variation (BCV) values across twelve diverse bulk RNA-Seq datasets. For each dataset, quantities inside the parentheses indicate their respective sample sizes. The blue bars indicate the BCV value for the first group, whereas the maroon bars denote the BCV value for the second group in each dataset. Group heteroscedasticity is determined by the difference between group-specific BCV values for each dataset. A greater difference in the BCV values between the groups, as also evidenced by a larger difference between the heights of the blue and maroon bars, indicates a higher degree of group heteroscedasticity within the respective datasets.

To this end, we propose a novel statistical framework under the robust linear model setup, denoted as Robseq (<u>Rob</u>ust Differential Expression Analysis of RNA-<u>seq</u> data). This framework effectively models diverse RNA-Seq data by considering heteroscedasticity and outliers in the estimation while maintaining robustness against FDR inflation across both small and large sample sizes. We validated our model by conducting extensive simulation studies, where we comprehensively benchmarked our proposed model against popular DE methods. Our findings underscore the superior performance of Robseq compared to the state-of-the-art DE tools in terms of FDR control, sensitivity, and specificity. Additionally, in two published population-level RNA-Seq datasets, we demonstrated that Robseq uncovers new biological insights that cannot be revealed by existing approaches. Finally, to facilitate practical applications, Robseq is publicly available as a user-friendly R package at https://github.com/schatterjee30/Robseq.

## 2 Results

### Corroborating the presence of group heteroscedasticity in RNA-Seq data

Previous studies have noted the presence of group heteroscedasticity in RNA-Seq data ^15,39,10,50,72^; however, the implementation of models that account for group heteroscedasticity remains scarce. This section aims to emphasize this phenomenon further by presenting additional evidence that group heteroscedasticity is not merely an incidental occurrence in RNA-Seq data but a common feature. We achieve this by analyzing bulk RNA-Seq datasets ranging from small- to large-scale RNA-Seq studies across ten diverse human diseases and two human tissues (**Table S1**). Strikingly, for each of these datasets, we demonstrate the presence of group heteroscedasticity (**Fig. 1**). Our approach mirrors that of a previous study ^72^ which identified group heteroscedasticity through higher biological coefficient of variation (BCV) values in one group compared to another. BCV, a key indicator of biological variability in gene expression among biological replicates, is widely used for estimating gene expression variance in RNA-Seq data ^45^. Following previous studies ^72^, here we demonstrate group heteroscedasticity by comparing the variability in group-specific BCV values and visualize this effect in the form of distinctive positions and shapes of group-specific mean-variance trend curves across these twelve diverse datasets (**Fig. S1**). We observe that the disparity in group-specific BCV values is significantly greater in large-scale RNA-Seq data, especially in those derived from the TCGA consortium studies ^62^. As an additional illustration of group heteroscedasticity, we also generated the p-value distributions for each dataset (**Fig. S2**), which did not align with the expected uniform distribution, implying the presence of unequal group variances. Combined, this evidence further highlights the pervasive nature of group heteroscedasticity in RNA-Seq datasets, emphasizing the critical need for robust statistical methods to account for this phenomenon in data analysis and interpretation.

### Robseq - A novel framework for statistically robust differential expression analysis

We have developed Robseq as a novel, flexible framework designed for conducting differential gene expression analysis of vastly different kinds of RNA-Seq data. This framework is compatible with both long and short read sequences and adeptly manages both equal and unequal biological variations across groups, known as group homoscedasticity and heteroscedasticity, respectively. Robseq comprises three main components: (i) a robust linear model (RLM) using Huber weights to estimate robust log-fold changes across experimental conditions, (ii) the MacKinnon-White variance estimator for robust standard error calculation under group heteroscedasticity, and (iii) a t-statistic with modified degrees of freedom to enhance the identification of DE genes while performing inference. Specifically, the first step in Robseq’s framework involves using the RLM model with Huber weights, which ensures robust estimation of log-fold changes while minimizing the influence of outliers. This linear modeling approach also accommodates biological covariates of interest (such as exposure and phenotype) as well as confounders (such as age and gender), which are often crucial to adjust for in RNA-Seq studies. In the second step, Robseq employs the MacKinnon-White variance estimator. This is especially pertinent for data exhibiting heteroscedasticity. It corrects the estimated standard error obtained in the first step and offers a heteroscedastic-consistent standard error estimate. Lastly, Robseq conducts per-gene tests to assess differential expression effects. This is done using a Welch t-statistic that incorporates modified degrees of freedom based on the Bell-McCaffrey correction ^30^, effectively balancing the type-I error rate and maintaining high statistical power in DE analysis. Robseq is available as an open-source R package at https://github.com/schatterjee30/Robseq.

### Simulation settings

We assessed six different models for analyzing differential expression in RNA-Seq data (**Methods**). Two of these methods were based on negative binomial (NB) assumptions, which include the widely employed DESeq2 ^40^ and edgeR.LRT (edgeR + likelihood-ratio test) ^53^, while two operate under a nonparametric setup, including dearseq ^21^ and the classical Wilcoxon rank-sum test. The remaining two methods involve two variants of the limma model for differential expression analysis of RNA-Seq data, referred to as voom ^33^ (limma + voom transformation) and voomByGroup ^72^ (limma + group-specific trends).

To evaluate these methods in diverse settings, including our proposed method Robseq, we generated synthetic RNA-Seq data using SimSeq, a nonparametric simulation engine ^6^. SimSeq uses an existing RNA-Seq dataset as template data to ensure that the simulated datasets capture the essential features of the original dataset such as count distributions and variability (**Methods**). Each simulated dataset comprised a total of 10, 000 genes, with the number of DE genes set to 1, 000. We focused on two sample sizes, *n* = 20, 50 per experimental group, to assess model performance in both small and large samples leading to four distinct scenarios: **Scenario 1**, featuring a small sample size data (*N* = 40) with homoscedastic template data; **Scenario 2**, involving a small sample size data (*N* = 40 with heteroscedastic template data; **Scenario 3**, including a large sample size data (*N* = 100 with homoscedastic template data; and **Scenario 4**, comprising a large sample size data (*N* = 100 with heteroscedastic template data. To ensure the robustness of our results and reduce the impact of random sampling errors, these simulation scenarios are replicated 100 times.

To generate synthetic data for these four scenarios, we supplied two different template datasets to SimSeq. Specifically, for the heteroscedastic scenarios, we supplied a colon cancer dataset ^36^ to SimSeq. This dataset comprised of 27,284 genes with measured expression profiles across non-tumor and tumor samples. Notably, these groups exhibited heteroscedasticity, with group-specific BCV values of 0.66 and 0.73 respectively (**Fig. 1**). In contrast, for the homoscedastic scenarios, we supplied a Chronic Pulmonary Obstructive Disease (COPD) dataset ^32^ to SimSeq. This dataset contained 24,782 genes, but unlike the colon cancer dataset, it demonstrated homoscedasticity in the expression profiles across healthy controls and COPD patients, with group-specific BCV values of 0.5.

Following the generation of synthetic data, we evaluated several key aspects for performance evaluation: i) The ability of the models to control type-I error rate at an imposed level of significance - this involved determining the proportion of genes labeled as DE at a 5% significance threshold by each model when there was no actual difference in mean gene expression between the groups; ii) The ability of the models to identify DE genes (sensitivity/power) - to assess the power of a model when true positive ground truth is known, we calculated the ratio of true positives identified by a respective model to the total number of actual positives; iii) The ability of the models to control FDR at the 5% nominal level - to assess the FDR of a model, we calculated the ratio of true positives identified by a respective model to the total number of actual positives; and iv) The prediction accuracy of the models - we evaluated this using Matthews Correlation Coefficient (MCC). MCC offers a balanced measure by taking into account all quadrants of the confusion matrix: true positives, true negatives, false positives, and false negatives, providing a comprehensive measure of a model’s performance. To assess the control of the type-I error rate, findings from all four scenarios are considered. However, for other evaluations such as FDR control, statistical power, and MCC, we primarily focused on results from Scenarios 2 and 4, which involved heteroscedastic data, while results from the other scenarios (Scenarios 1 and 3) are in the **Supplementary Materials**. Likewise, for the analysis of real data, we followed the same approach, documenting the population-level DE analysis of a heteroscedastic dataset in the main article and the DE analysis of a homoscedastic dataset in the **Supplementary Materials**.

### Robseq controls false positives when no differentially expressed genes are present

In RNA-Seq studies, assessing the type-I error (false positive rate) when testing the null hypothesis is crucial to ensure the reliability of DE detection. This assessment measures the likelihood of incorrectly classifying genes as differentially expressed. Due to the complex variability in RNA-Seq data, it is essential to control the type-I error to prevent the spread of erroneous findings, thereby maintaining the biological significance and reproducibility of the results. In the context of this evaluation, when synthetic data contains no truly DE genes, the p-values for individual genes should align with a uniform distribution, a phenomenon known as well-calibrated p-values. A deviation from this pattern would indicate the presence of false positives, questioning the validity of the model’s findings. In other words, effective models should maintain the type-I error rate such that the proportion of p-values remains below a specific significance threshold.

We observed that across both sample sizes (N = 40, 100) and under homoscedastic conditions (Scenarios 1 and 3), all models effectively controlled the type-I error rate at the nominal 5% level (**Fig. 2**). Specifically, when N = 40, the Wilcoxon test, voom, and voomByGroup were noteworthily conservative in their approach. Meanwhile, dearseq and Robseq exhibited a moderate level of conservatism. On the other hand, DESeq2 and edgeR.LRT managed to uphold the type-I error rate at the 5% threshold. When the sample size was increased to N = 100 in the homoscedastic scenario, the Wilcoxon test, voom, and voomByGroup continued to demonstrate conservative behavior and dearseq remained moderately conservative, whereas Robseq, DESeq2, and edgeR.LRT sustained the type-I error rate at the 5% mark. Conversely, in Scenarios 2 and 4, which involved heteroscedastic conditions for both sample sizes, the NB-based methods, specifically DESeq2 and edgeR.LRT, tended to be overly liberal, with their type-I error rates exceeding the 5% threshold. In contrast, the nonparametric methods (dearseq, Wilcoxon) and the linear models (Robseq, voom, voomByGroup) consistently maintained proper control over the type-I error rate in these heteroscedastic scenarios. In these contexts, voom and voomByGroup were quite conservative, whereas dearseq, Wilcoxon test, and Robseq were moderately conservative. Additionally, we evaluated the models at a more stringent 1% level. The results of this evaluation closely mirrored those observed at the 5% level (**Fig. S3**) for all the models except for DESeq2 and edgeR.LRT which failed to control their type-I error rates at the 1% level in all of the scenarios. Furthermore, to determine whether our results varied with a significantly larger sample size, we carried out a similar assessment using an expanded sample size of N = 140. We found that, for all the models, the results were generally in line with those acquired for the sample size of N = 100. However, in this instance, both DESeq2 and edgeR.LRT did not maintain control of their type-I error rates at the 1% and 5% thresholds across all scenarios. (**Fig. S4**).

**Figure 2:**
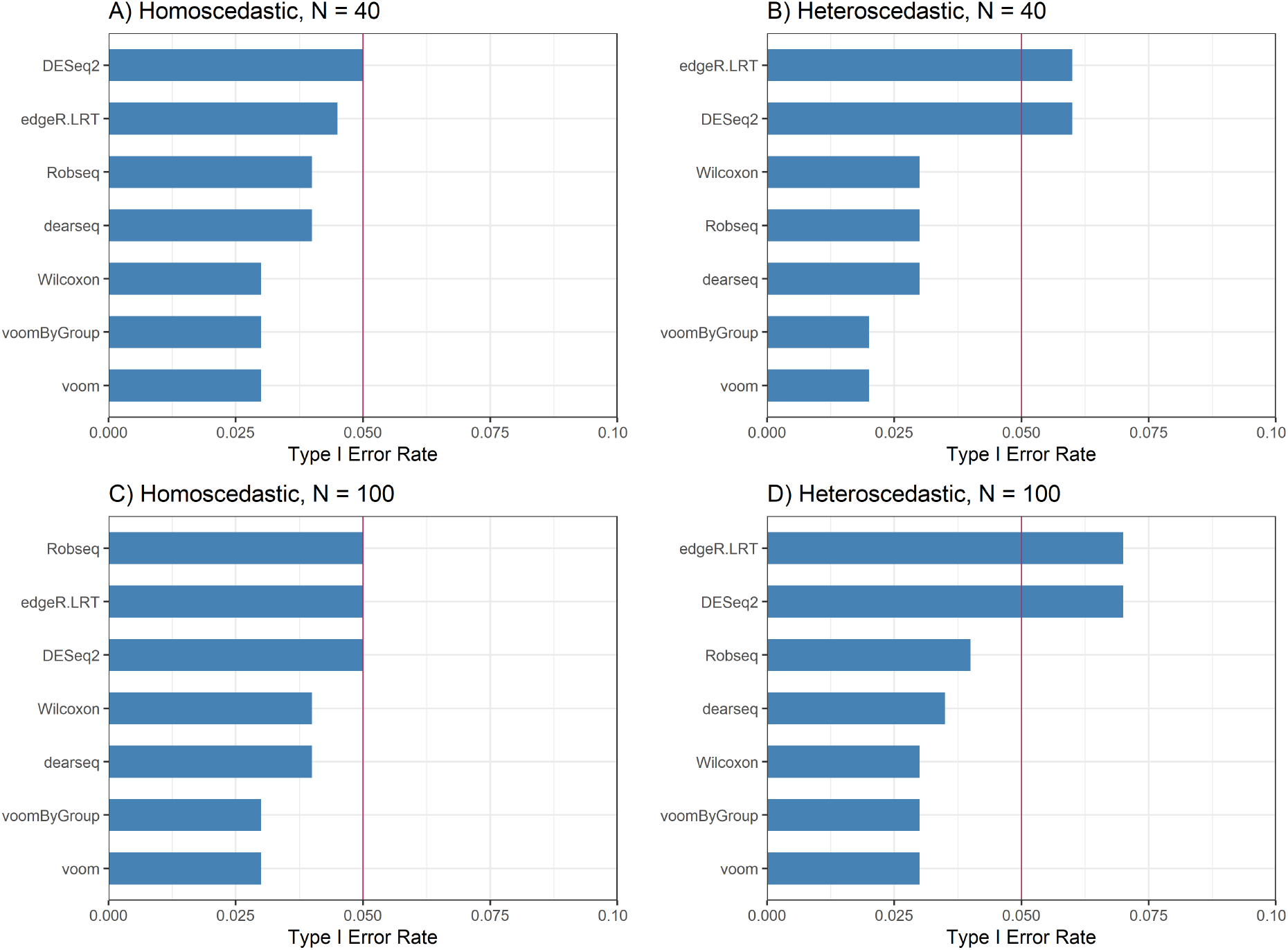
Presence of heteroscedasticity can obscure type-I error rate control for popular DE methods when the true differential expression is absent. The bar plots illustrate the percentage of genes that have an unadjusted p-value less than 0.05 for each method in four scenarios: (**A**) when data is homoscedastic with a sample size N = 40, (**B**) when data is heteroscedastic with a sample size N = 40, (**C**) when data is homoscedastic with a sample size N = 100, and (**D**) when data is heteroscedastic with a sample size N = 100. A red line in these plots represents the nominal type-I error rate, set at 0.05. These results are compiled from the median of 100 simulations. Methods that effectively regulate the type-I error at or below this nominal threshold will have their corresponding bars positioned beneath the red line

### Robseq has good control over false discovery rate while maintaining substantial power

Following the evaluation of type-I error control in scenarios without DE genes, we focused on assessing the statistical power and FDR control of various models for accurately detecting true differences in gene expression at a 5% significance level. Assessing FDR is vital because it gauges the reliability and validity of findings in significant gene expression, ensuring that these reflect genuine biological changes rather than mere random fluctuations. On the other hand, assessing a model’s statistical power is essential to verify its effectiveness in identifying genes with significant expression changes. This is particularly important in functional genomics experiments where differential expression effects are modest and difficult to detect.

In the context of FDR control, **Fig. 3** illustrates the FDR of various models at a 5% significance threshold, with 10% of genes marked as DE under heteroscedastic conditions. In these cases, for all sample sizes, all models except edgeR.LRT and DESeq2 managed to maintain FDR control at the 5% level. The Wilcoxon test, voom, and voomByGroup consistently showed conservatism across different sample sizes, while Robseq displayed moderate conservatism. Interestingly, dearseq displayed moderate conservatism under the sample size of N = 40 while was highly conservative under the sample size of N = 100. Noticeably, edgeR.LRT and DESeq2 consistently exhibited inflated FDRs, reinforcing previous study findings ^55,12,38,21,54^. Similarly, in homoscedastic scenarios (**Fig. 5**), most models effectively controlled the FDR for both the sample sizes, except for edgeR.LRT and DESeq2. Here again, the Wilcoxon test, voom, and voomByGroup was conservative, while dearseq and Robseq showed moderate conservatism for both sample sizes.

**Figure 3:**
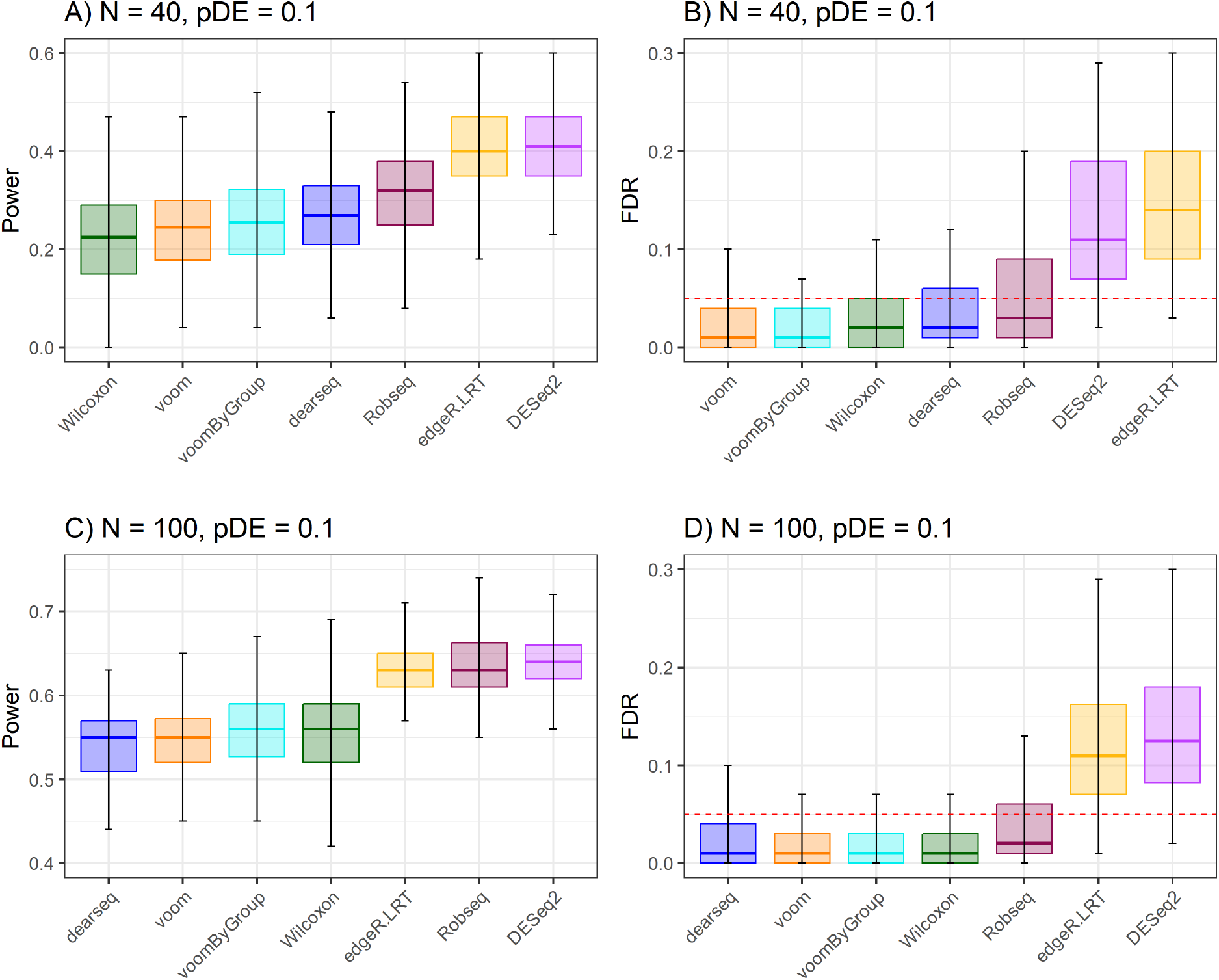
The impact of group heteroscedasticity on statistical power and FDR inflation in synthetic bulk RNA-Seq data. We simulated 100 synthetic data under a heteroscedastic scenario with two different sample sizes and with the true proportion of DE genes (pDE) being 10%. Panel (**A**) and (**C**) show the boxplot of statistical power from 7 DE methods arranged left to right in increasing values of median sensitivity for *N* = 40 and *N* = 100, and panel (**B**) and (**D**) show the FDR for the same simulation experiments. Methods corresponding to the boxplots on the right side of the power plots are generally more efficient in finding DE genes. Methods that effectively manage to keep the FDR at or below the predetermined nominal level (0.05) should be positioned below the red dotted line.

Regarding statistical power, first we report the percentage of true positives identified by different models based on a Benjamini-Hochberg adjusted p-value threshold of 5% under heteroscedastic conditions (**Fig. 3A-C**). When N = 40, Robseq demonstrated the highest power among methods that previously showed control over type-I error rate and FDR. Although edgeR.LRT and DESeq2 showed higher statistical power than Robseq, they were not considered due to their inability to control the type-I error rate and FDR. Particularly, when sample sizes increased to N = 100, Robseq showed the highest overall power among all the models compared. Remarkably, the power difference between Robseq and the model with the second highest power that also controlled FDR (Wilcoxon test) was around 10%. In the homoscedastic scenario (**Fig. S5A-C**), with a sample size of N = 40, Robseq had the second highest power among models that controlled type-I error rate and FDR, with dearseq having the highest power. However, as the sample size increased to N = 100, Robseq emerged as the most powerful model among those controlling FDR and type-I error rate. In addition, considering that the proportion of DE genes might affect the performance of DE models ^12^, we evaluated the models in both heteroscedastic and homoscedastic scenarios, and for both sample sizes, with 20% of genes labeled as DE (**Figs. S6-S7**). Like the scenario with 10% DE genes, we found with 20% DE genes Robseq stood out to be the model with the highest power. Furthermore, to check if the model performances changed with a much higher sample size, we evaluated the models under the heteroscedastic scenario with a sample size of N = 140 (**Fig. S8**). We found that Robseq had the highest power in cases with either 10% or 20% DE genes. Specifically, Robseq’s power (73%) was 7% higher than the Wilcoxon test (66%), which had the second-highest power when the DE genes proportion was 10%. Similarly, when the DE genes proportion was 20%, Robseq’s power (77%) remained 5% higher than the Wilcoxon test (72%). Overall, these findings indicate that Robseq is highly effective in detecting true signals while also maintaining robust control over FDR.

### Robseq has superior performance according to Matthews Correlation Coefficient

The Matthews Correlation Coefficient (MCC) is an important metric for assessing a model’s performance in accurately classifying genes as DE or not in differential gene expression studies. It adeptly balances the rates of true positives, true negatives, false positives, and false negatives, crucial for the correct identification of DE genes and for minimizing classification errors in non-differentially expressed genes. MCC’s thorough evaluation method is key to preventing significant interpretive errors in biology by effectively addressing both false negatives and positives. This makes MCC a robust and dependable metric, thereby enhancing the validity and credibility of outcomes in gene expression research. MCC scores range from -1 to 1, with higher scores such as 1 corresponding to a perfect classifier, while a negative MCC indicates a strong disagreement between prediction and observation. In our evaluation (**Fig. 4**), with a sample size of N = 40 under the heteroscedastic scenario and 10% DE genes, Robseq ranked second highest in MCC among all models, whereas for a sample size of N = 100, Robseq achieved the highest MCC score. When evaluating scenarios with 20% DE genes under heteroscedastic conditions, Robseq consistently outperformed other models in MCC scores for both sample sizes. Additionally, under homoscedastic conditions, Robseq demonstrated superior MCC scores compared to other models for most of the scenarios (**Fig. S9**). We also evaluated Robseq’s performance with a sample size of N = 140 under heteroscedastic conditions and found that it maintained the highest MCC score (**Fig. S10**). These results indicate that Robseq exhibits superior accuracy in correctly classifying genes as either DE or not DE in our simulation studies.

**Figure 4:**
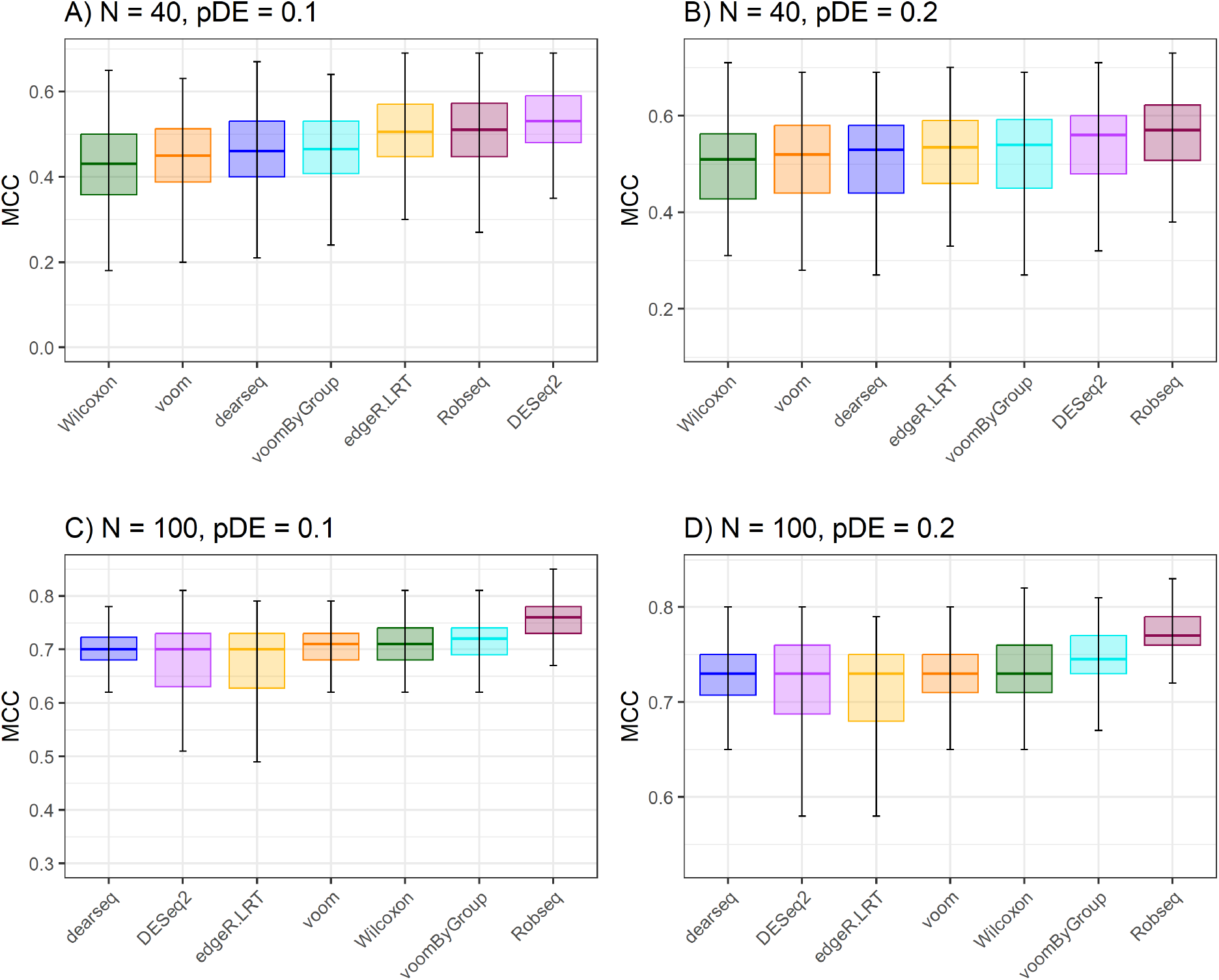
Robseq has superior MCC scores compared to existing models. We simulated 100 synthetic data under a heteroscedastic scenario with two different sample sizes and two levels of proportions of DE genes (pDE). Panel (**A**) and (**C**) show the boxplot of MCC scores from 7 DE methods when the true proportion of DE genes in the simulated dataset is 10%, and, (**B**) and (**D**) show the MCC scores of the simulation experiments with the true proportion of DE genes being 20%. The boxplots are arranged in the increasing order of the median MCC scores from left to right in each plot. Methods with high MCC scores more accurately classify the genes in DE and non-DE categories.

**Figure 5:**
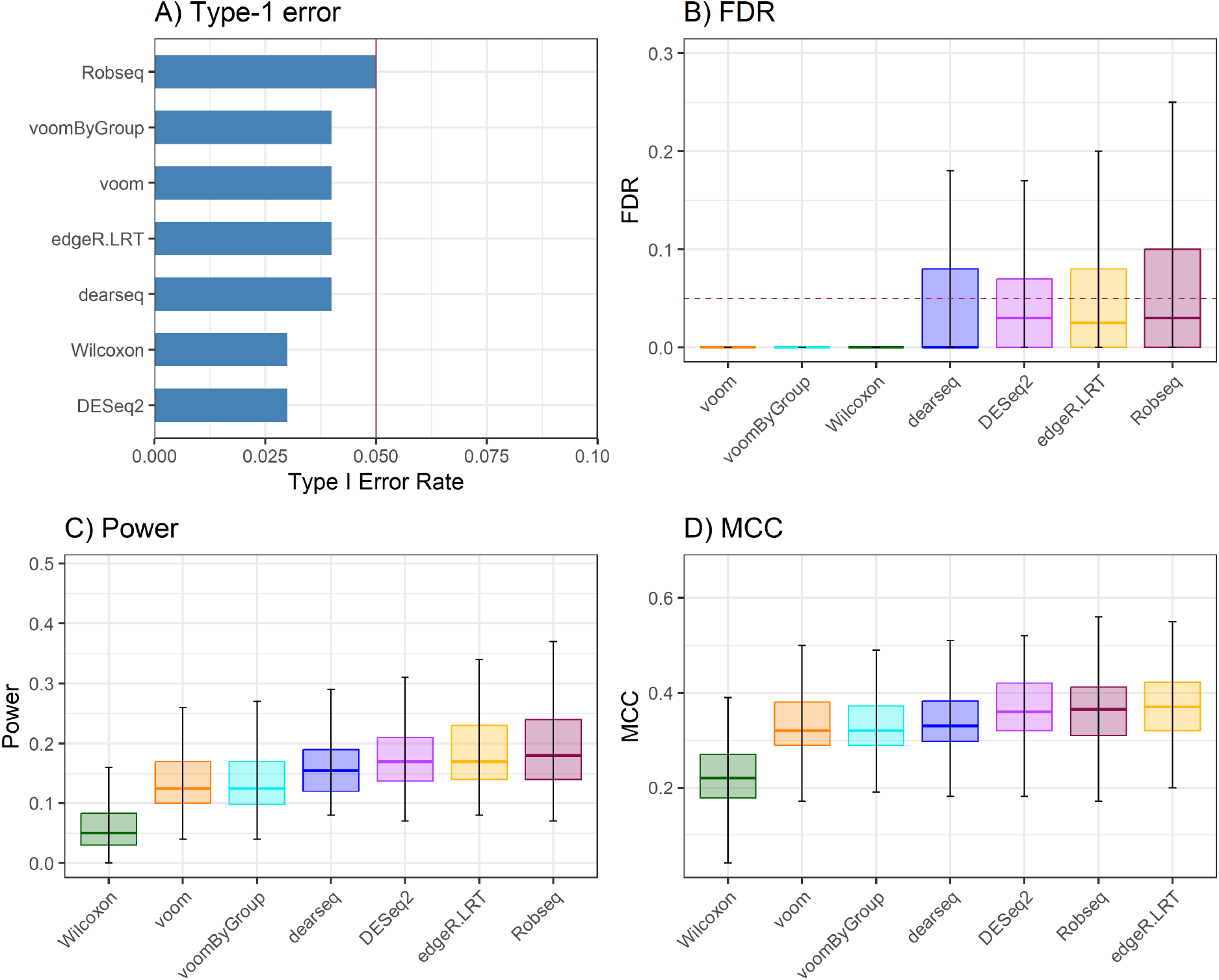
Robseq has superior performance in modeling long non-coding RNA expression data. We simulated 100 synthetic data using a long non-coding RNAseq dataset as a template where the total sample size is N = 30 and the proportion of DE genes is fixed at 10%. The boxes in each boxplot are arranged according to the increasing value of median metrics from left to right. Methods that effectively maintain the type-I error rate at the predetermined nominal level are expected to be positioned to the left of the vertical red line which represents the specified nominal type-I error rate of 5% (**A**). Most methods effectively manage to keep the FDR at or below the predetermined nominal level (0.05) in this dataset as they are all positioned below the red dotted line (**B**). Methods with higher power (**C**) and MCC (**D**) are better at identifying both DE and non-DE genes.

### Robseq offers additional utility by maintaining favorable performance in modeling long non-coding RNA expression profiles

The exploration and examination of long non-coding RNA (lncRNA) holds significant importance in understanding human health and diseases due to their extensive yet largely unexplored functional role within the genome. Various studies on gene expression have revealed that lncRNAs are typically expressed at low levels, exhibit high levels of variability, and demonstrate tissue-specific expression patterns ^66,64,51^. These traits pose considerable challenges for DE tools and could potentially impact their effectiveness adversely ^2^. In light of this, we wanted to additionally assess the effectiveness of Robseq in detecting DE in lncRNA gene expression.

In this simulation study, we utilized a Psoriasis dataset as our reference data, focusing specifically on long non-coding RNAs (lncRNAs). We narrowed down the dataset to include only long intergenic non-coding and antisense RNAs, resulting in 2959 lncRNAs measured across two groups: healthy controls and Psoriasis patients, totaling N = 52 samples. Most lncRNAs exhibited low expression levels and high intra-group variability, consistent with previous research. We also observed significant variability between groups, with a between-group coefficient of variation (BCV) of 0.4 for healthy controls and 0.56 for Psoriasis patients, indicating group heteroscedasticity. To generate synthetic count data, we employed SimSeq using the lncRNA expression profiles from the Psoriasis dataset. Since the subset of lncRNAs in the original dataset was less than 10,000 genes, we augmented it to this size by sampling with replacement. Each simulated dataset comprised 10,000 genes, with the number of differentially expressed (DE) genes set to either 1,000 (10%) or 2,000 (20%). We focused on a sample size of n = 15 per experimental group, resulting in two distinct scenarios: Scenario 1 with N = 30 and 10% DE genes, and Scenario 2 with N = 30 and 20% DE genes. To ensure robustness and reduce the impact of random sampling errors, each simulation scenario was replicated 100 times.

In our evaluation of both scenarios, we first examined how well various models controlled the type-I error rate at a 5% significance level and found that all models effectively managed to control it (Fig. 5A). We also assessed the control of the type-I error rate at a more stringent 1% level (Fig. S11A), and here too, all models demonstrated effective control. Secondly, regarding false discovery rate (FDR) control, we observed that with both 10% and 20% DE genes in the data (Figs. 5B, S11B), all models maintained FDR control at the 5% significance level. However, models like voom, voomByGroup, and Wilcoxon were more conservative in FDR control compared to edgeR.LRT, DESeq2, dearseq, and Robseq. Thirdly, in terms of statistical power, Robseq showed the highest power, followed by edgeR.LRT, in scenarios with both 10% and 20% DE genes (Figs. 5C, S11C). Lastly, concerning Matthews Correlation Coefficient (MCC) scores, Robseq again scored the highest, closely followed by edgeR.LRT, in scenarios with both 10% and 20% DE genes (Figs. 5D, S11D). These results suggest that Robseq performs substantially well, even in challenging scenarios such as the DE analysis of lncRNAs. This additional assessment underscores Robseq’s versatility in modeling different RNA-Seq biotypes while maintaining superior performance.

### Robseq uniquely deciphers potentially mechanistic insights from published population-level bulk RNA-Seq studies

Recognizing that simulations may not fully reflect the complexity of real biological systems due to their reliance on simplified assumptions, we next analyzed two publicly available real-world, large-scale bulk RNA-Seq datasets that depict group heteroscedasticity. This allowed us to test Robseq’s ability to identify DE genes in population-level bulk RNA-Seq datasets, thus bridging the gap between theoretical simulations and practical applications. We applied Robseq in conjunction with four representative methods (voom, voomByGroup, Wilcoxon, and dearseq) that demonstrated FDR control in our simulations, which recapitulated the superior performance of Robseq observed in simulations compared to established gold standard methods.

The first dataset examined in our analysis was a Lupus dataset ^47^, comprising 24,408 gene expression profiles across 120 samples after quality control. Lupus, also known as Systemic Lupus Erythematosus (SLE), is a chronic inflammatory autoimmune disease that affects various organ systems, posing challenges for diagnosis. Common symptoms include skin rashes, malar rash, arthritis, pleurisy, serositis, alopecia, and Lupus nephritis ^1^. The causes of Lupus are not fully understood, but it is believed to be influenced by a combination of genetic and environmental factors, which may trigger or exacerbate symptoms. At the cellular level, the disease process initiates with interactions between the adaptive and innate immune systems, leading to elevated cytokine levels, complement activation, immune complex deposition, and subsequent inflammation and tissue damage ^1^. Among other characteristics, this dataset exhibited group heteroscedasticity (**Fig. 1**), as evidenced by a BCV of 0.29 in the healthy group compared to 0.36 in the SLE group. Previous analyses of this dataset did not account for this inherent group heteroscedasticity, prompting us to re-analyze it using Robseq to potentially glean better biological insights.

Robseq identified a total of 8,266 DE genes, the highest number compared to all other models. Additionally, there was a significant overlap of these genes (*N* = 5, 322) with those identified by other models (**Fig. 6**). Conspicuously, Robseq uniquely detected 381 DE genes not identified by other models (**Table S2, Fig. 7B**). The gene ontology (GO) over-representation analysis showed that the DE genes overlapping with other models were primarily involved in RNA processing and modifications, immune response signaling, and responses to viral infections (**Fig. S12A**). In contrast, the DE genes uniquely identified by Robseq were associated with pathways related to the immune effector process, platelet formation, DNA damage and repair, and hormonal pathways influencing maternal processes (**Fig. 7A**).

**Figure 6:**
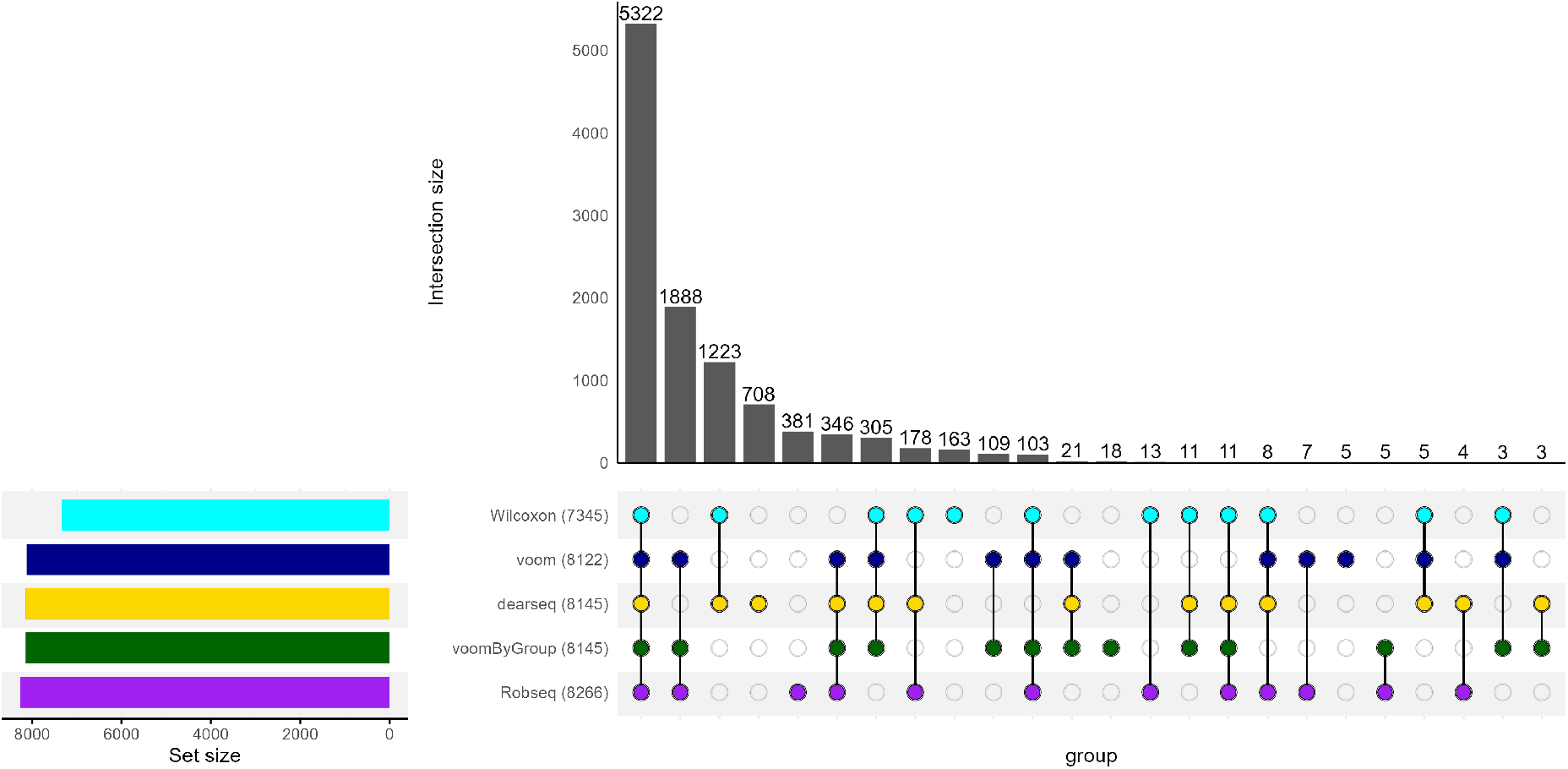
UpSet plot of number of DE genes detected across five DE models in the Lupus data. Using five DE models, we summarize the number of DE genes (out of 24,408 genes) detected between healthy control (N = 58) and systemic Lupus erythematosus (N = 62) groups in the Lupus dataset. Numbers in parentheses represent the total number of DE genes identified by the corresponding method. Genes with observed FDR smaller than *α* = 0.05 were deemed as DE.

**Figure 7:**
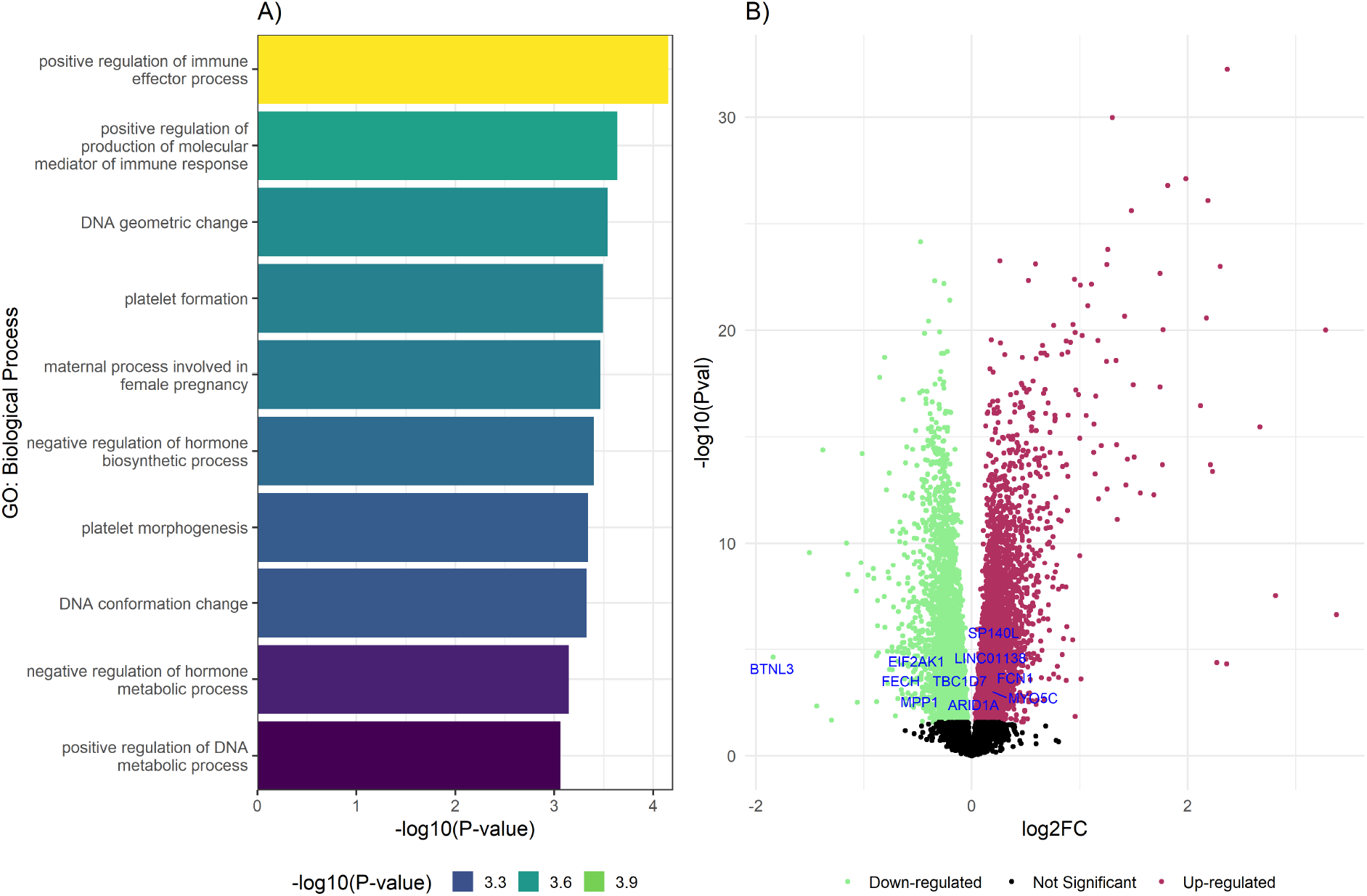
Immune and DNA-repair pathway genes are involved in Lupus which were uniquely identified by Robseq. In panel **(A)**, the bars represent the top ten enriched GO biological process terms associated with DE genes that were uniquely identified using Robseq. The x-axis of the chart shows the -log10 transformed (unadjusted) p-values for the top 10 GO biological process terms. Additionally, a color scale is used to group the strength of the association based on the p-values. In Panel **(B)**, a volcano plot illustrates the DE analysis of the Lupus dataset conducted with Robseq. The x-axis of this plot displays the per-gene log_2_ fold changes in the Lupus dataset, while the y-axis shows the -log10 transformed p-values for these genes. The points in green signify genes identified as DE with downregulation, whereas the points in red indicate genes recognized as DE with upregulation. The points in black represent genes that were not DE in the Lupus dataset. The top ten DE genes, chosen for having the lowest FDR-adjusted p-values and uniquely identified by Robseq, are highlighted with blue text.

One of the top hits uniquely identified by Robseq in the Lupus data was the gene Butyrophilin-like protein 3 (BTNL3), which was found to be downregulated in patients with SLE compared to healthy controls (Table S2). BTNL3 acts as a co-stimulatory molecule regulating human and mouse *γδ* T cell subsets ^31,27^. These *γδ* T lymphocytes, along with B cells and *αβ* T cells, form a crucial part of the adaptive immune system in all vertebrates ^70^. Prior studies have shown that BTNL3 binds to Vg4+ T-cell receptors in a manner akin to superantigens ^70^. Crucially, there is evidence that BTNL3’s role in selecting and maintaining *γδ* T cells is significant in curbing the progression of complex intestinal diseases that cause tissue damage ^14^. This finding is particularly relevant, given that many of the common risk alleles associated with genetic predisposition to SLE are found in genes related to the immune system. Consequently, Robseq’s identification of BTNL3 as DE underscores the importance of recognizing a significant and previously unlinked gene, allowing us to identify novel pathways related to Lupus (**Table S2, Fig. 7B**). Additionally, increasing evidence suggests that abnormal DNA repair plays a part in the progression of Lupus. DNA repair and cell cycle control protein TP53BP1 and DNA-associated modification and editing factors such as XRCC5, EXO5, APOBEC3F, and APOBEC3G were uniquely identified by Robseq, which not only were missed by other algorithms but also were not previously known to associate with Lupus. Furthermore, epigenetic regulatory factors such as ARID1A, DNMT3A, and histone H3.3 chaperone HIRA were uniquely identified as dysregulated genes in Lupus by Robseq. Molecular function analysis for the Robseq-identified unique genes significantly associated with DNA-binding and epigenetic regulation. These findings collectively highlight that Robseq was able to identify biologically relevant unique DE genes which were previously unknown to be linked with SLE.

The second dataset we examined was a colon cancer dataset, featuring 27,284 gene expression profiles across 144 samples after quality control. Colorectal cancer (CRC) is one of the most common cancers worldwide, known for its high morbidity and mortality. Research indicates that CRC is genetically driven, involving alterations in numerous oncogenes and tumor suppressor genes ^7^. Several genes and proteins related to CRC have been identified, playing key roles in cell proliferation, differentiation, apoptosis, and metastasis ^28,61^. However, the precise molecular mechanisms of CRC remain elusive. This dataset showed significant intra-group variability and group heteroscedasticity, with a BCV of 0.66 in the non-tumor group compared to 0.73 in the tumor group. To our knowledge, no previous study has performed DE analysis on this dataset, providing a unique opportunity to potentially obtain novel DE signals associated with CRC using Robseq, aiming for enhanced biological understanding.

Robseq identified a total of 11,417 DE genes, the highest number compared to all other models. Similar to the Lupus data analysis, there was a significant overlap of Robseq-identified DE genes (8,391 in total) with those identified by other models (**Fig. 8**). Robseq uniquely detected 452 DE genes that were not identified by other models (**Table S3, Fig. 9B**). The GO analysis showed that the DE genes overlapping with other models were primarily involved in ribosome biogenesis, cytoplasmic translation, and mitochondrial gene expression (**Fig. 12B**). In contrast, the DE genes uniquely identified by Robseq were associated with pathways related to subcellular transport, adhesion, and energy metabolism (**Fig. 9A**). Among the notable genes uniquely identified by Robseq in the CRC dataset is the solute carrier family 10 member 5 (SLC10A5), which was found to be underexpressed in tumor samples compared to non-tumor samples (**Table S3**). SLC10A5, is a bile acid transporter primarily found in the liver and kidneys, has been identified as being downregulated in colon cancer ^20^. Another notable gene uniquely identified by Robseq was the gene exocyst complex component 8 (EXOC8). This gene, part of the exocyst complex crucial for positioning exocytic vesicles at fusion sites on the plasma membrane ^13^, also showed reduced expression in tumor samples (**Table S3**). Interestingly, many downregulated genes identified by Robseq are associated with phosphatidylinositol, actin, dynein, myosin, and spectrin binding (**Table S3**). In particular, APC, a cell polarity regulator and a tumor suppressor, is mutated in familial adenomatous polyposis and colorectal cancer ^74^. APC regulates both actin and microtubule cytoskeletons, forming the central machinery for cell migration ^19^. These findings collectively highlight that genes uniquely identified by Robseq indicate the potential involvement of skeletal proteins in colon cancer which other models failed to identify. These genes could serve as potential new therapeutic targets for effectively managing colon cancer; however, extensive research is needed to identify their clinical relevance to colon cancer.

**Figure 8:**
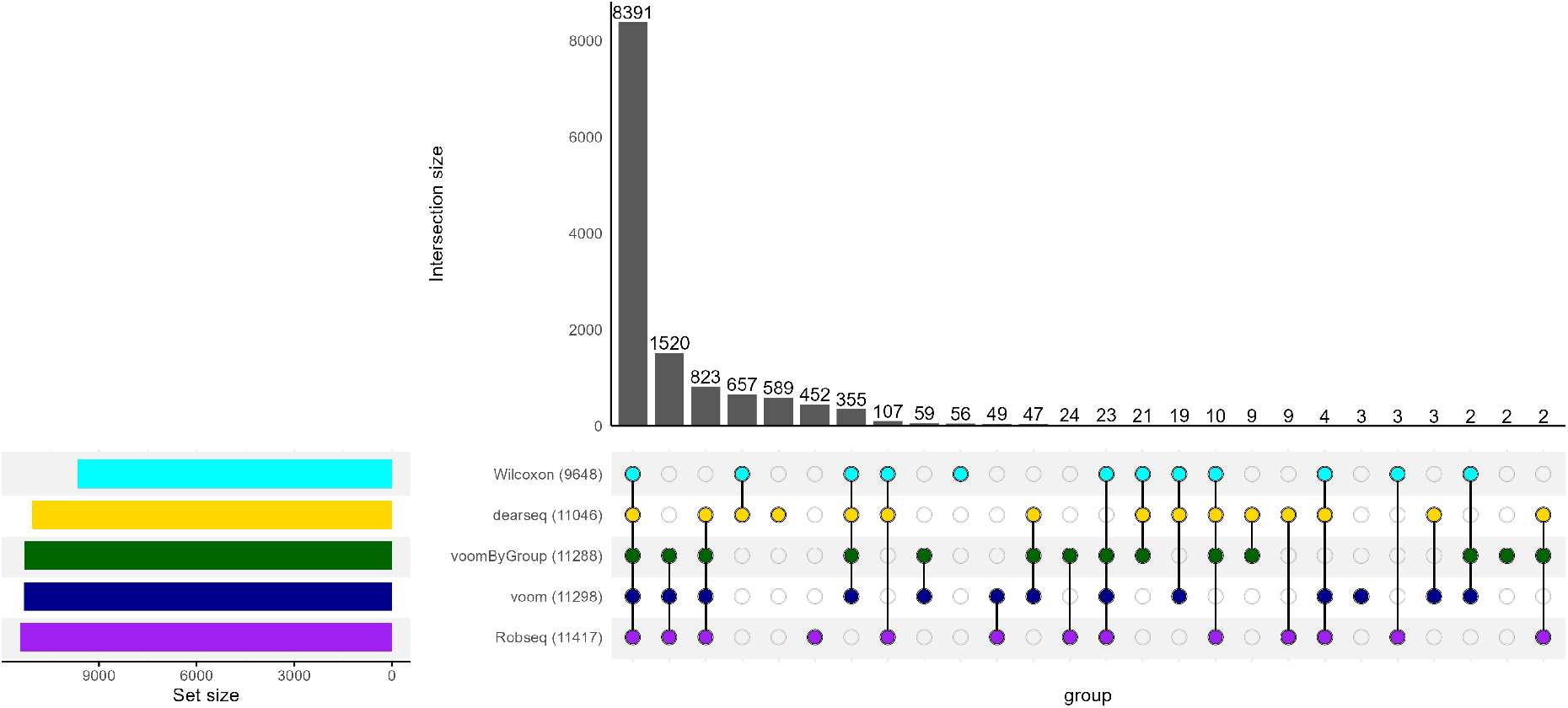
UpSet plot of number of DE genes detected across five DE models in the colon cancer data. Using five DE models, we summarize the number of DE genes (out of 27,284 genes) detected between non-tumor (N = 72) and tumor samples (N = 72) in the colon cancer dataset. Numbers in parentheses represent the total number of DE genes identified by the corresponding method. Genes with observed FDR smaller than *α* = 0.05 were deemed as DE.

**Figure 9:**
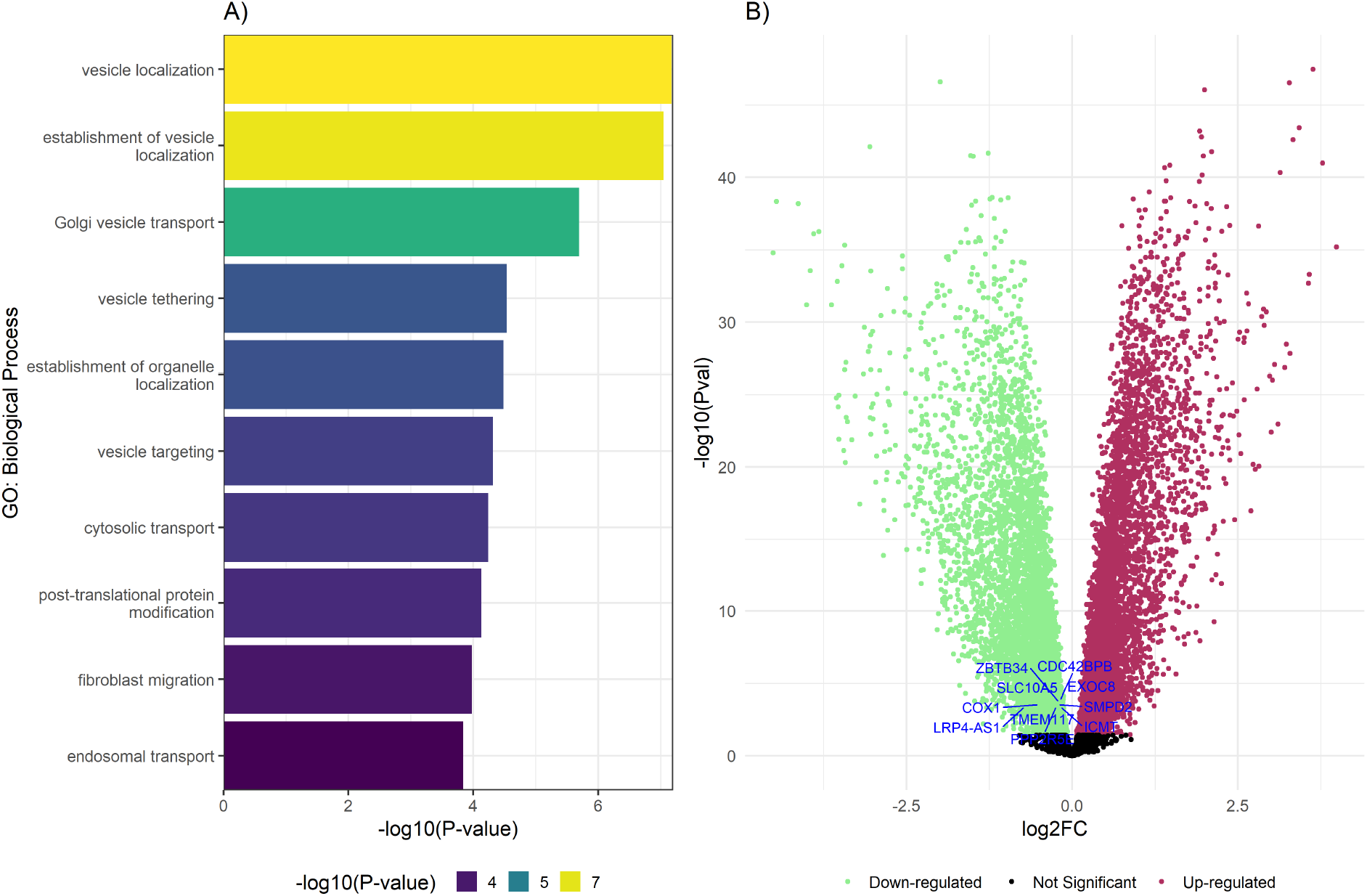
Subcellular transport and energy metabolism pathway genes are involved in colon cancer which were uniquely identified by Robseq. In panel **(A)**, the bars represent the top ten enriched GO biological process terms associated with DE genes that were uniquely identified using Robseq. The x-axis of the chart shows the -log10 transformed (unadjusted) p-values for the top 10 GO biological process terms. Additionally, a color scale is used to group the strength of the association based on the p-values. In Panel **(B)**, a volcano plot illustrates the DE analysis of the colon cancer dataset conducted with Robseq. The x-axis of this plot displays the per-gene log_2_ fold changes in the colon cancer dataset, while the y-axis shows the -log10 transformed p-values for these genes. The points in green signify genes identified as DE with downregulation, whereas the points in red indicate genes recognized as DE with upregulation. The points in black represent genes that were not DE in the colon cancer dataset. The top ten DE genes, chosen for having the lowest FDR-adjusted p-values and uniquely identified by Robseq, are highlighted with blue text.

As additional illustrations, we examined two more datasets (**Supplementary Materials**). The first, a Chronic Obstructive Pulmonary Disease (COPD) dataset ^32^, features 24,345 genes, measured across 189 samples post-quality control. We verified that this dataset demonstrates group homoscedasticity, indicated by an identical BCV of 0.5 in both healthy and COPD groups. Corresponding to results from our homoscedastic simulation study, this population-level re-analysis corroborated that Robseq detected the highest number of DE genes, including several COPD-associated genes uniquely found by Robseq (**Supplementary Materials**). The second additional dataset on Psoriasis ^23^ encompasses 2959 lncRNAs with expression levels measured across 52 samples following quality control. This dataset showed group heteroscedasticity, with a BCV of 0.4 in healthy samples and 0.56 in the Psoriasis group. This re-analysis paralleled our lncRNA simulation study findings, depicting that Robseq detected the highest number of DE genes, including several Psoriasis-associated genes uniquely discovered by Robseq. These re-evaluations and additional analyses confirm Robseq’s superior performance in detecting DE genes across diverse RNA-Seq datasets, showcasing its efficacy in capturing biologically relevant signals in both heteroscedastic and homoscedastic data, as well as its ability to identify DE genes across diverse RNA biotypes.

## 3 Discussion

Maintaining well-calibrated p-values and reliable false discovery control is crucial in gene expression studies, or in any high-throughput study for that matter where multiple testing is involved, ensuring reproducible, trustworthy, and actionable findings that are truly indicative of biological changes, rather than merely random variations. Consequently, FDR control at the desired level upholds the scientific integrity of past and present gene expression research and differential expression analyses, as failure to control the FDR gives researchers a false sense of certainty about the reliability of their discoveries. Despite the availability of several high-profile DE methods spanning more than a decade of research, controversy persists due to recent evidence showcasing a complete failure of popular state-of-the-art DE methods to control the FDR at the nominal level. Similar tendencies have been prominent in other related nascent fields such as microbiome and single-cell differential analysis ^24,43,44^. However, the extent to which this phenomenon has been observed in a mature field like gene expression and bulk RNA-Seq is especially noteworthy.

In this study, we hypothesized that group heteroscedasticity (unequal variability in gene expressions among experimental groups) is a common aspect in RNA-Seq data, which may contribute to the increase in false discoveries observed in gold standard models such as edgeR ^53^ and DESeq2 ^40^. This aspect has been largely overlooked in the literature, but we show strong evidence that group heteroscedasticity is a common occurrence in RNA-Seq data rather than an isolated event. As one of the key highlights of this study, we provide comprehensive evidence that group heteroscedasticity may pose a potential risk for generating false positives in the results of several gold standard models. Our findings also reveal that models that do control false discoveries in heteroscedastic conditions are often underpowered. To overcome these limitations in published methods, we have introduced a new method, Robseq, which essentially bridges the gap between recall and power. Robseq utilizes a heteroscedasticity-consistent statistical pipeline, offering a robust and powerful approach for DE analysis. It effectively controls the FDR at the nominal level while maintaining superior power — a challenge not fully addressed by current methods.

To demonstrate Robseq’s effectiveness as a flexible and powerful DE method, we conducted extensive simulation studies across a variety of data settings under a nonparametric setup, which is a crucial consideration as strong parameter assumptions can violate the reliability of benchmarking studies, as indicated in the literature ^24,38^. These diverse settings ranged from generating homoscedastic to heteroscedastic synthetic data, encompassing both small and large sample sizes and including different proportions of true DE genes. Notably, our choice to employ a nonparametric approach for simulations ensured that the evaluation of models was conducted impartially, without favoring any specific parametric models, such as those based on an NB distribution, thereby guaranteeing fair and unbiased comparisons between statistical models. Additionally, we assessed Robseq’s versatility by evaluating its performance in lncRNA data analysis, a biotype of RNA-Seq data characterized by low abundance and high noise levels, which leads to subpar performance when published RNA-Seq DE analysis tools are applied without modifications to this data type. Our results demonstrated that regardless of (i) sample size, (ii) number of true DE genes, (iii) RNA-Seq biotype, and (iv) heteroscedasticity (or lack thereof), Robseq consistently managed to control the FDR across these diverse use cases while achieving significantly higher statistical power compared to published DE methods. Furthermore, Robseq showed high accuracy in classifying both DE and non-DE genes, as evidenced by its strong MCC scores in most simulation scenarios. Among the three gold standard models we compared (DESeq2, edgeR, and limma-voom), edgeR and DESeq2 often struggled with FDR control, aligning with recent findings in the literature^38^. This issue might stem from non-uniform distributions of p-values generated by these models, a phenomenon found in previous studies and reflected in our simulations. It’s important to note that this might not be a general characteristic of these models, as indicated by our lncRNA simulation results, where all models, including edgeR and DESeq2, effectively controlled false discoveries. These findings suggest that the most popular RNA-Seq DE models may not always ensure accurate control of false positives or exhibit the highest statistical sensitivity in various RNA-Seq study settings, and Robseq presents a flexible and powerful alternative mitigating these issues across experimental designs and data types.

In addition to the superior performance demonstrated by Robseq in our comprehensive simulation studies, we also reanalyzed three population-scale bulk RNA-Seq datasets and one lncRNA dataset, which highlights Robseq’s effectiveness in identifying biologically significant transcriptomic signals with relevance to disease outcomes. Particularly, the DE analysis of complex diseases such as Lupus, COPD, colon cancer, and psoriasis using Robseq revealed its superior ability not only in detecting the highest number of DE genes from these datasets but also in uncovering novel, unique DE genes with biological importance that other models failed to identify. For example, the results of the GO over-representation analysis demonstrated that the unique DE genes identified by Robseq were associated with a range of biological processes, including immune response, DNA damage and repair, epigenetic regulation, cytoskeletal function, and hormonal pathways. These processes, not identified by other DE models, are crucial in understanding the complexity of these diseases. Further, Robseq’s identification of the gene BTNL3 as DE in the Lupus data has underscored the potential significance of previously unlinked genes in the disease. Similarly, the detection of genes such as TP53BP1, XRCC5, EXO5, APOBEC3F, APOBEC3G, ARID1A, DNMT3A, and HIRA, which were missed by other models, shed light on the roles of DNA repair, cell cycle control, and epigenetic regulation in Lupus, which may lead to enhanced understanding of the molecular mechanisms of this disease. Additionally, Robseq’s discovery of various cytoskeletal genes that were differentially regulated in colon cancer, which other models did not detect, is particularly noteworthy. This finding is significant as the role of cytoskeletal proteins in cancer development is often underemphasized, yet Robseq’s results suggest that these genes could have potential implications in colon cancer. Furthermore, in the study of psoriasis, Robseq identified 80 unique non-coding DE genes, including several novel lncRNAs with previously unknown functions. Collectively, these findings emphasize Robseq’s capability to identify unique genes that are differentially regulated and linked to biological processes not recognized by other models. This highlights its utility in discovering potential new links between previously unrelated genes, paving the way for identifying potential new biomarkers for various complex diseases.

From a practical standpoint, Robseq offers a comprehensive and general solution for gene-level analysis of vastly different kinds of RNA-Seq data. For example, Robseq does not require the input data to be strictly counts (i.e., integers) as would be the case in methods like DESeq2, which rely on a negative binomial distribution assumption. Given that many Bioinformatics pipelines return RNA-Seq data in terms of normalized counts, widely available in databases such as GEO, the flexibility of Robseq stands out as it accommodates all these variations in its robust, platform-agnostic linear model framework. Embedding Robseq in a robust, heteroscedasticity-aware linear model framework further allows it to be highly flexible for conducting gene-by-gene analysis of RNA-Seq gene expression data under diverse experimental settings. The implementation of Robseq is modular, allowing the robust linear model to be paired with a variety of preprocessing and normalization methods. This underscores the adaptability and versatility of Robseq, a significant departure from published methods, many of which require the use of specific normalization methods to generate results.

The application of Robseq beyond bulk RNA-Seq profiles, such as single-cell data (both at the cell-level and pseudo-bulk level), remains a significant avenue for future research. One notable constraint in such extensions is the efficient handling of an excessive amount of zeroes and the additional variability posed by single-cell technologies ^57,75^. Integrating methodologies like empirical Bayes shrinkage and Bayesian hierarchical approaches could offer valuable solutions ^52^, given their track record of providing stable estimates in noisy settings. Cell type-specific DE analysis, where multiple cell types are present, requires further modification, as it necessitates the incorporation of cell type compositions from heterogeneous tissues ^46^. Additionally, the increasing availability of RNA-Seq studies with spatial and temporal characteristics means that Robseq’s robust linear model must be adapted to accommodate these complex settings. Finally, while Robseq is highly scalable to population-level data, as demonstrated by the real data applications of this study, leveraging low-level programming ^18^ and distributed computing ^48^ could enhance its scalability further, which we hope to explore as a future direction. Combined, such extensions will allow researchers to analyze diverse, large-scale RNA-Seq datasets, moving towards a unified estimation framework accommodating a variety of experimental designs and biotypes. We believe that the Robseq framework and its improved DE detection performance across varied scenarios represent an important step in this direction, which can ultimately aid in discovering consistent biological signals and carefully targeted follow-up experiments, reducing costly and time-consuming experimental efforts in current and future mechanistic studies based on RNA-Seq profiles.

## 4 Methods

### Modeling Strategy

Let 𝒢 be the total number of observed genes and 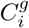 be the raw count of the *g*^*th*^ gene for the *i*^*th*^ sample (*i* = 1,…, *N*). To build our model, we rely on the linear model setup for each gene *g* where expression values 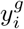 for each of these genes have been log_2_ normalized. Specifically, our model is;

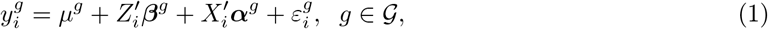

where 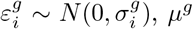 is the intercept, ***β***^*g*^ represents the effect of DEA identifiers in *Z*_*i*_. *X*_*i*_ contains additional covariates/confounders that the count 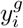 depends on. The superscript *g* in eq. (1) identifies individual models for each gene, and within each model, the errors are independently distributed. A statistically significant non-zero ***β***^*g*^ indicates that gene *g* is DE.

Unique characteristics of typical RNA-Seq data along with the above model specification pose two challenges, viz., (i) the effect of small sample size and outliers on the estimation of regression parameters and (ii) handling the heteroscedastic *σ*_*i*_. An ordinary least square estimate is highly susceptible to both the presence of an outlier ^4^ and unequal variance ^56^. Hence, we opt for robust estimation of the model parameters via robust linear model (RLM) regression. Moreover, we also use a bias-adjusted variance estimator in our proposed test statistic for hypothesis testing.

### Estimation of robust regression coefficients

To safeguard against the biased estimation of model parameters due to not satisfying the assumptions in the linear model, *RLM* ^29^ is used to regress 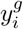 in eq. (1). The basic idea of *RLM* is to use weights to modify the influence a data point can have on model parameters. The weights along with the model parameters, are estimated in an iterative scheme commonly known as iterative re-weighted least square (IRLS).

In particular, we use the RLM implementation in the MASS package ^65^ in R software ^49^ that uses *M* - estimation ^4^ method to estimate (***β***^*g*^, ***α***^*g*^). With *M*-estimation, the estimate 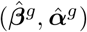 is determined by minimizing the following objective function,

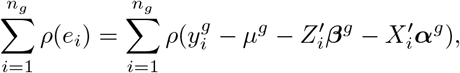

where the function *ρ*(·) provides the contribution of each residual to the objective function. Minimizing the above objective is equivalent to solving the following system of equations:

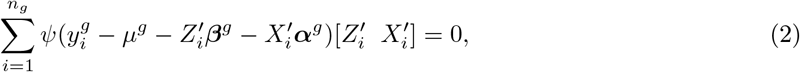

where *ψ* = *ρ*^*′*^, the derivative of *ρ*. By defining the weights *ω*(*e*) = *ψ*(*e*)*/e*, the above estimation is the same as solving a weighted least square problem. Huber estimator ^29^ is opted for the choice of *ρ*, in which the objective function and the corresponding weights are defined as follows:

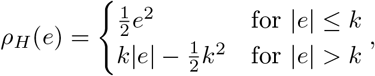

and

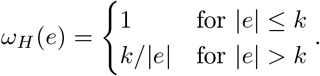

*k* serves as a tuning constant; smaller values of *k* produce more resistance to outliers but at the expense of lower efficiency when the errors are normally distributed, which is rarely the case. Finally, solving eq. (2) produces a robust estimate of ***β***^*g*^ and ***α***^*g*^, which can be expressed as:

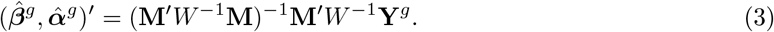

*W* denotes the matrix whose diagonals are the estimated weights *ω*, **M** = [***Z* X**] is the combined model matrix and **Y**^*g*^ is the vector of 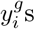.

### Inference under heteroscedasticity with general regressors

Concerning this study’s context, we have a linear regression setup with multiple regressors. For the sake of brevity, we will denote the full vector of regression parameters by ***β***^*g*^ and the full model matrix by **X** hereafter instead of (***β***^*g*^, ***α***^*g*^) and [***Z* X**] respectively. While RLM can provide a robust estimate of the regressors’ coefficients, however, the variance estimate of 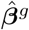 is generally biased. There exist commonly used estimators for 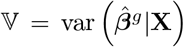 under several model assumptions. For instance, one of the first and most popular proposed estimators of variance under a heteroscedastic setup was by Eicker-Huber-White (EHW) for the single binary regressor case. Despite its applicability, later studies found the EHW variance estimator to be downward biased in the case of finite samples. Following that, MacKinnon and White proposed the HC2 variance estimator, which improved on the downward bias problem of the EHW estimator. It was later shown that the HC2 estimator, in general, removes only part of the finite sample bias in the EHW estimator, but in the case of a single binary regressor, the HC2 correction removes the entire bias. The 𝕍_HC2_ estimator under the general regressor setup has the following form:

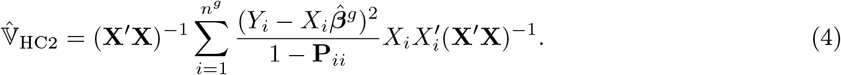

where **P** = **X**(**X**^*′*^**X**)^−1^**X**^*′*^ is the *N ×N* projection matrix, with the *i*th column **P**_*i*_ = **X**(**X**^*′*^**X**)^−1^*X*_*i*_ and (*i, i*)th element denoted by 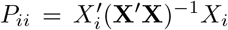. Let Ω be the *N×N* diagonal matrix with the *i*th diagonal element equal to *σ*^2^(*X*_*i*_), and let *e*_*N,i*_ be the *N*-vector with the *i*th element equal to 1 and all other elements equal to 0. Let *I*_*N*_ be the *N × N* identity matrix. The residuals 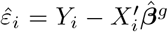 can be written as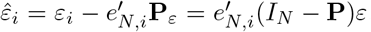, or in vector form,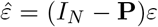. For more details on the variance estimators for single binary regressor case, see ^5^.

### Hypothesis testing

According to the proposed model in eq. (1), a gene *g* is DE if ***β***^*g*^ is significantly different from 0. Thus, Robseq tests for the null hypothesis for each gene *g*:

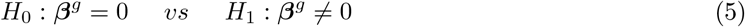

Following the variance estimate discussed above, a t-statistic for the *k*th component of ***β***^*g*^ can written as 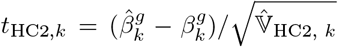. 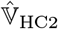 is unbiased for 𝕍, however, the sampling distribution of the *t* statistic is not normally distributed. Whereas in the homoscedastic case, the *t*_HC2,*k*_-statistic has an exact *t*-distribution. Welch (1951) proposed an approach to deal with the problem of variance estimation of ***β***_*g*_ under heteroscedasticity (also known as the Behrens-Fisher problem), which is to approximate the distribution of the *t*-statistic by a *t*-distribution with degrees of freedom adjustment that reflects the variability of the variance estimator 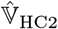. Welch proposed a constant factor *K* such that the distribution of 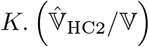 had a chi-square distribution with degrees of freedom equal to *K*. As, 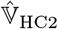 is independent of 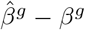 under normality, 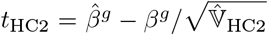 would attain a *t*-distribution with degrees of freedom *K*. As there exists no such *K* for which 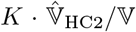 follows an exact chi-squared distribution, Welch suggested approximating the scaled distribution of 𝕍_HC2_ by a chi-squared distribution, with the degrees of freedom parameter *K* chosen such that 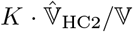 has the first two moments in common with a chi-squared distribution with degree of freedom equals to *K*. Following the same principle, Bell and McCaffrey ^5^ (BM hereafter) proposed the following solution to determining *K*. Under homoscedasticity, 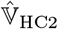 is unbiased, leaving us to solve for *K* by equating only the second moments for those two distributions. Under normality, 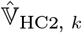 is a linear combination of *N* independent chi-squared 1 random variable,

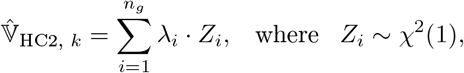

where all *Z*_*i*_s are independent and the weights *λ*_*i*_ are the eigenvalues of the *N × N* matrix of *σ*^2^ *·* **G**^*′*^**G**, with the *i*th column of the *N × N* matrix **G**, is equal to

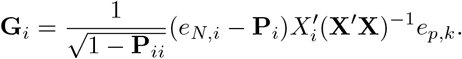

Given these weights, the BM degrees of freedom that match the first two moments of [ineq to that of a chi-squared *K* distribution is given by

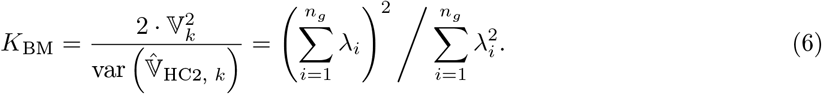

The p-values were obtained from the 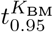 distribution in a gene-specific manner. Furthermore, an adjustment (the Benjamini–Hochberg procedure) was applied on p-values to account for multiple-testing. DE status of a gene was inferred based on its adjusted p-value falling below the 5% significance level.

### Datasets

Publicly available bulk RNA-Seq datasets that were analyzed in this article include:

- Lupus data - Blood samples from SLE patients and healthy donors. This dataset is accessible from the Gene Expression Omnibus (GEO) database under accession number GSE112087.
- COPD data - Lung tissue samples from subjects with normal spirometry and COPD patients. This dataset is accessible from the GEO database under accession number GSE57148.
- Colon cancer data - Colon tissue samples from tumor and adjacent normal tissue. This dataset is accessible from the GEO database under accession number GSE156451.
- Psoriasis data - Lesional skin samples from Psoriasis patients and normal skin samples from healthy individuals. This dataset is accessible from the GEO database under accession number GSE74697.

Additional publicly available bulk RNA-Seq datasets that were examined in **Fig. 1** are included in **Table S1**.

### Data normalization, filtering, and enrichment analysis

Before conducting DE analysis, we filtered genes with low expression/abundance levels using the *filterByExpr* function from edgeR ^53^, with the exception of simulations and re-analyses involving lncRNA data. For the lncRNA data, given the focus on genes with low abundance, we only filtered out genes that exhibited no expression in all samples. Subsequently, normalization for each dataset was performed using the relative log expression (RLE) method using the *estimateSizeFactors* function in DESeq2 ^40^. To conduct GO analyses on the DE results from the four population-level RNA-Seq datasets (Lupus, CRC, COPD, and psoriasis), we used the *enrichGO* function in clusterProfiler (version 4.8.3), utilizing the org.Hs.eg.db annotations ^73^

### Synthetic data generation for simulation studies

We generated synthetic data for our simulation studies using an existing bulk RNA-Seq data simulator, SimSeq ^6^. Specifically, SimSeq is a nonparametric simulator designed to generate RNA-Seq read count matrices by sub-sampling from an existing RNA-Seq dataset. This simulation process begins with a representative template dataset, normalized using the TMM method from the edgeR package ^53^, with each gene being expressed in at least one sample from the groups being studied. The selection of DE genes is conducted through probability sampling without replacement, with the weighting inversely related to each gene’s local FDR. This rate is determined from the p-values of the Wilcoxon Rank Sum test, giving priority to genes more likely to be DE. Subsequently, a set of equivalently expressed (EE) genes is chosen using equal probability sampling from those not in the DE set. For these EE genes, a predetermined number of samples are selected from the source dataset. The number of samples selected from the source dataset is based on a minimum threshold to ensure balanced representation from both groups, thereby ensuring a comprehensive and representative gene analysis. For more information about SimSeq, refer to the original publication ^6^.

To determine the suitability of the synthetic data generated by SimSeq, we conducted several evaluations focusing on basic characteristics. This involved verifying whether the simulator, when provided with homoscedastic template data, produced data that maintained homoscedasticity (**Fig. S17**). Likewise, we examined whether heteroscedastic template data led to the creation of synthetic data that displayed heteroscedasticity (**Fig. S18**). Beyond these basic assessments, we also evaluated the synthetic data’s quality using a range of additional metrics suggested by Soneson and Robinson ^58^ (**Figs. S19-S20**). Overall, based on these assessments we found that the synthetic data generated by Simseq mostly conformed with the input template data as well as with the specific simulation scenarios.

### Identification of DE genes for competing models

All six competing models (DESeq2, edgeR.LRT, voom, voomByGroup, dearseq, and the Wilcoxon rank-sum test) took a read count matrix and a condition label vector as input. The parameters were set based on the user guides of these method’s software packages.

- For DESeq2 (v1.40.2), differential analysis was conducted using the *DESeq* function, followed by the extraction of results via the *results* function.
- For edgeR.LRT (v4.0.12), genes with low counts were first removed using *filterByExpr*, then normalized using the Trimmed Mean of M values (TMM) approach and finally, the likelihood ratio test was employed to identify DE genes.
- For voom (limma) (v3.58.1), we used a similar gene filtering and normalization as in edgeR. After this, the *voom* transformation was applied to the normalized, filtered count matrix, and differential analysis was carried out using *lmFit* and *eBayes* functions.
- For voomByGroup (limma), we again used a similar gene filtering and normalization as in edgeR. After this, the *voomByGroup* function was applied to the normalized, filtered count matrix, and differential analysis was carried out using *lmFit, contrasts*.*fit and treat* functions.
- For dearseq (v1.12.1), the initial steps of filtering and normalization mirrored those in edgeR. This was followed by the use of *dear* _*seq* function with an asymptotic test for detecting DE genes.
- The Wilcoxon rank-sum test also followed the same preliminary steps as edgeR. Here, we calculated p-values by inputting counts-per-million (CPM) values for each gene into the ‘wilcox.test’ function in R (v4.2.1), with a p-value threshold set based on the FDR using the BH method.

## Code Availability

Results were generated using R version 4.2.1. Robseq is available on https://github.com/schatterjee30/Robseq as an open-source software. Additionally, code for simulations, figures, and comparisons of DE analysis methods on the various datasets explored are also available at https://github.com/schatterjee30/Robseq.

## Acknowledgments

This research was supported in part by Lilly Endowment, Inc., through its support for the Indiana University Pervasive Technology Institute.

## Notes

https://github.com/schatterjee30/Robseq

### Competing Interest Statement

The authors have declared no competing interest.

