## Supplementary Materials for "Group Heteroscedasticity - A Silent Saboteur of Power and False Discovery in RNA-Seq Differential Expression"

### 1 Supplementary Figures

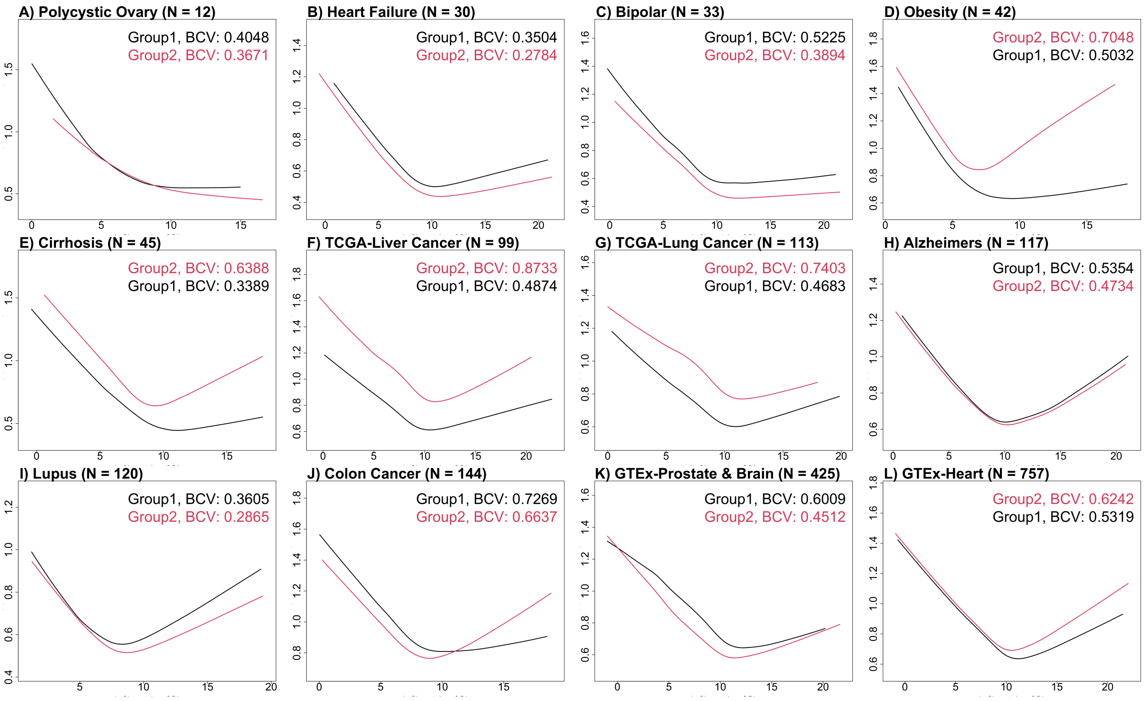

**Supplementary Figure** 1: **Illustration of group heteroscedasticity observed in bulk RNA-Seq data using mean-variance plots.** This plot illustrates the mean-variance trend curves specific to each group across twelve distinct bulk RNA-Seq datasets related to human diseases and tissues. Within each dataset, the sample sizes are noted in parentheses. The red curves represent the mean-variance trend of the first group, while the black curves represent the mean-variance trend of the second group in each dataset. The unique positions and shapes of these group-specific mean-variance trend curves are used to assess group heteroscedasticity, especially when comparing different groups within the same RNA-Seq dataset. The curve's shape and height indicate the overall variation within groups, encompassing both technical and biological factors. In datasets exhibiting group homoscedasticity, the mean-variance trend curves of the groups will be closely aligned and largely overlap. In contrast, datasets demonstrating group heteroscedasticity will show group-specific mean-variance trend curves that are distinct and mostly non-overlapping. Additionally, the numbers located in the top right corner of the plot area represent the group-specific biological coefficient of variation (BCV) values for each dataset.

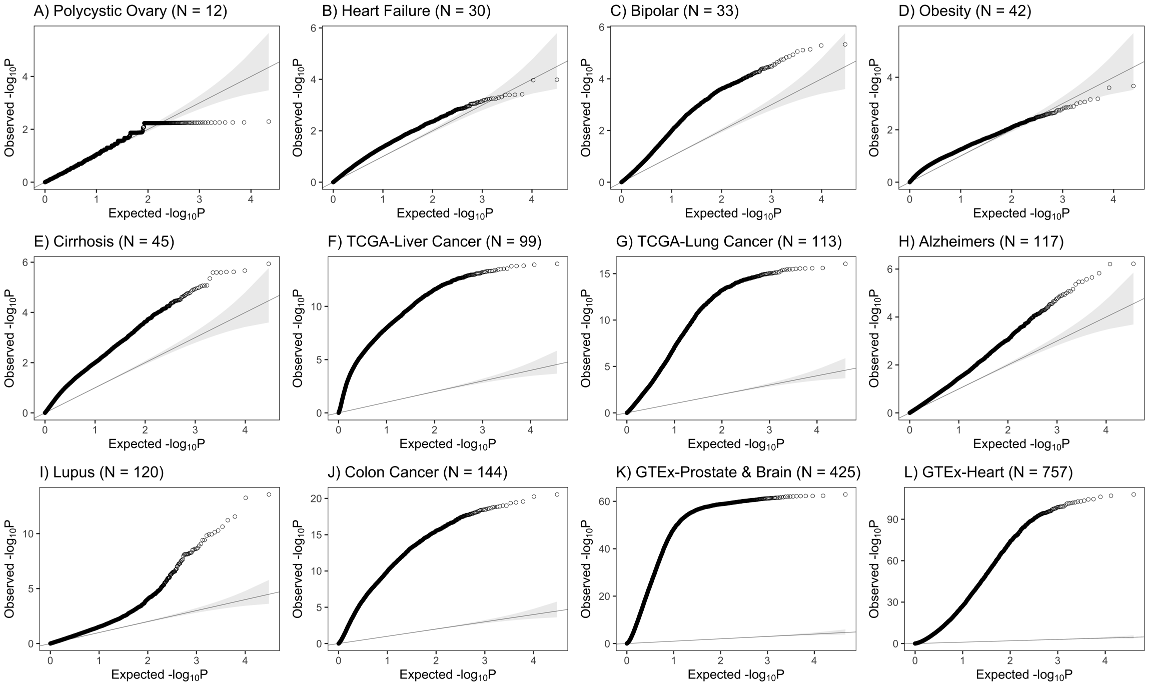

**Supplementary Figure** 2: **Illustration of group heteroscedasticity observed in bulk RNA-Seq data using statistical tests for homoscedasticity.** This graph presents Quantile-Quantile (QQ) plots for twelve distinct bulk RNA-Seq datasets related to human diseases and tissues. In each plot, the x-axis represents the theoretically expected p-values, while the y-axis shows the p-values observed in the data. These p-values originate from a statistical test that compares the observed between group variance to a uniform distribution, under the assumption that, in cases of homoscedasticity (equal variance across groups), the p-values should uniformly distribute. Essentially, if there is no evidence of group heteroscedasticity (varying variance across groups), the p-values would typically align with this uniform distribution. Marked departures from this expected uniform pattern are indicative of a violation of the null hypothesis, which assumes equal variance, thus suggesting the presence of group heteroscedasticity. The QQ-plots clearly demonstrate deviations from the expected uniform line, indicating the existence of group heteroscedasticity in these datasets.

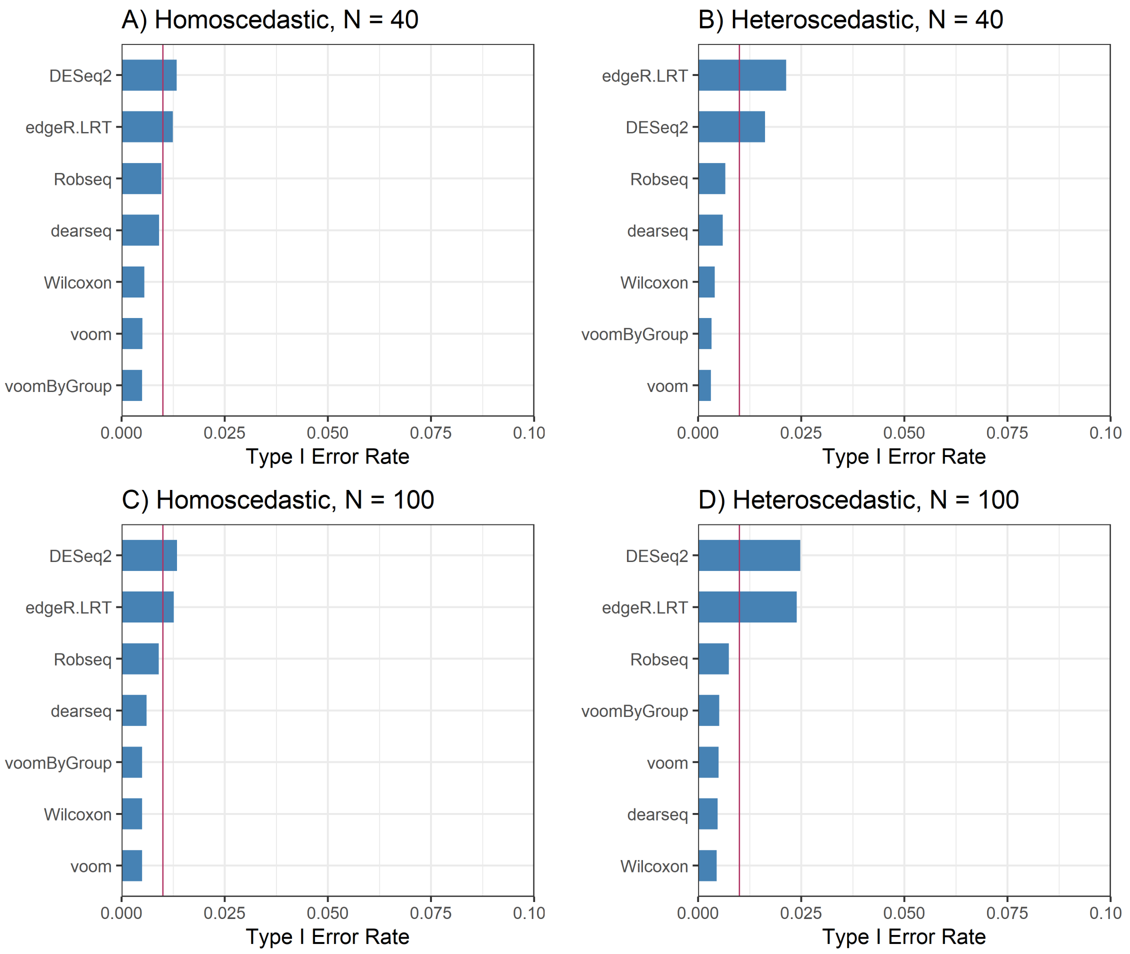

**Supplementary Figure** 3: **Type-1 error rates when true differential expression is absent.** The bar plots illustrate the percentage of genes that have a p-value less than 0.01 for each method in four scenarios: A) when data is homoscedastic with a sample size N = 40, B) when data is heteroscedastic with a sample size N = 40, C) when data is homoscedastic with a sample size N = 100 and D) when data is heteroscedastic with a sample size N = 100. A red line in these plots represents the nominal type I error rate, set at 0.01. These results are compiled from the median of 100 simulations. Methods that effectively regulate the type-1 error at or below this nominal threshold will have their corresponding bars positioned beneath the red line.

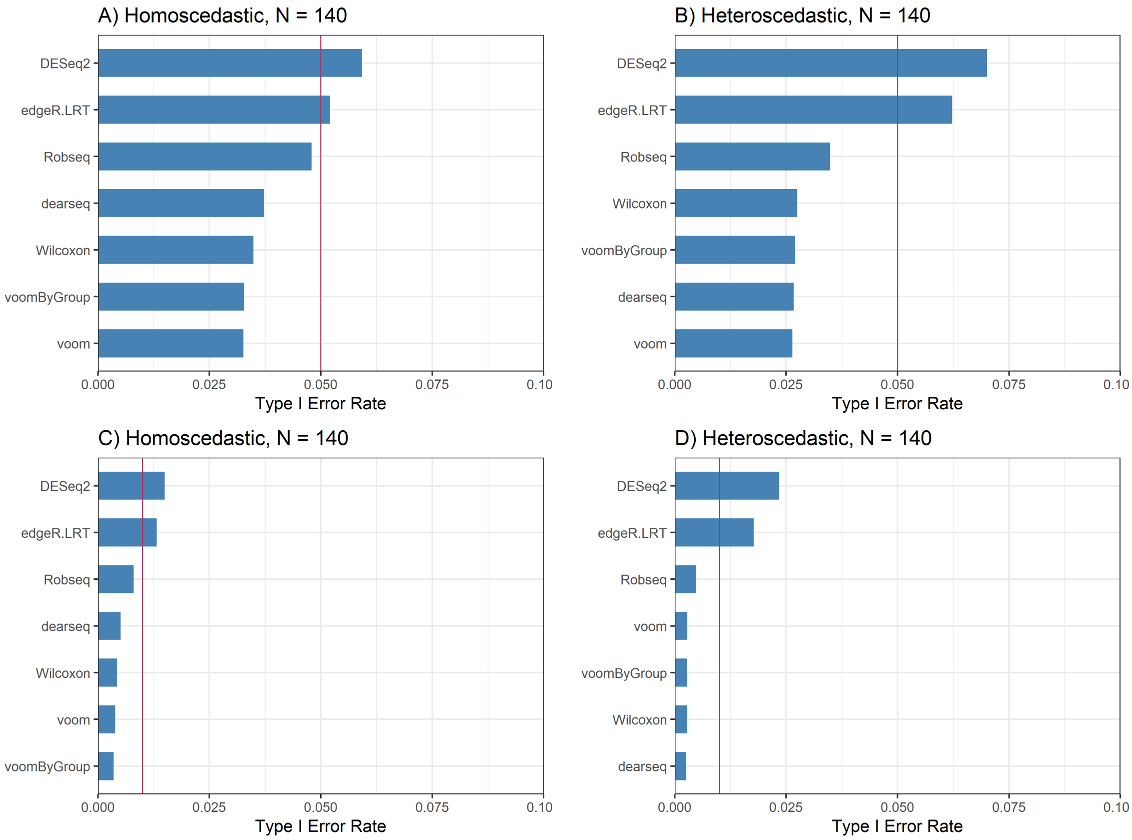

**Supplementary Figure** 4: **Type-1 error rates when true differential expression is absent.** The bar plots illustrate the percentage of genes that have a p-value less than a predetermined nominal threshold for each method in four scenarios: A) when data is homoscedastic with a sample size N $=$ 140 and p-value cut-off of $\alpha = 0.05$ , B) when data is heteroscedastic with a sample size N $=$ 140 and p-value cut-off of $\alpha = 0.05$, C) when data is homoscedastic with a sample size N $=$ 140 and p-value cut-off of $\alpha = 0.01$ and D) when data is heteroscedastic with a sample size N $=$ 140 and p-value cut-off of $\alpha = 0.01$. A red line in these plots represents the nominal type I error rate. These results are compiled from the median of 100 simulations. Methods that effectively regulate the type-1 error at or below this nominal threshold will have their corresponding bars positioned beneath the red line.

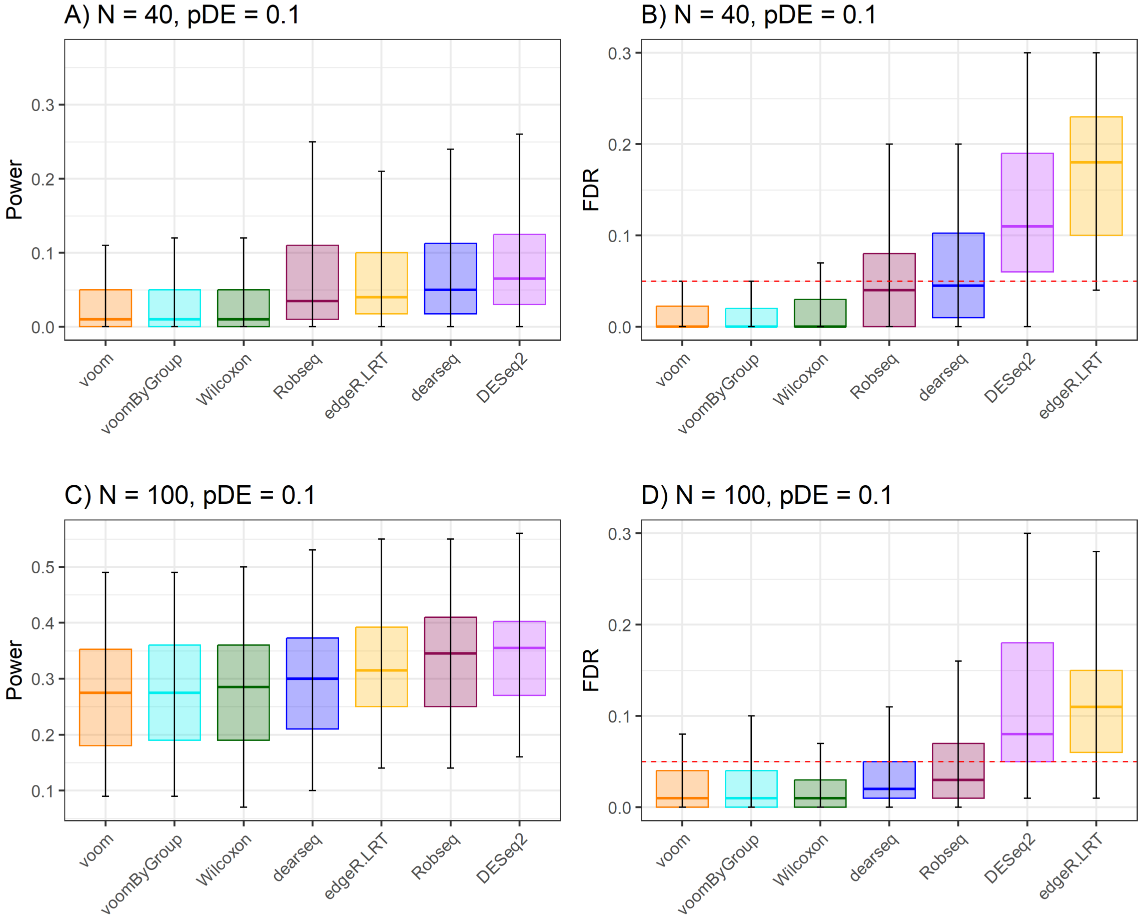

**Supplementary Figure** 5: **Power and FDR evaluations in homoscedastic data when 10% genes were labeled as differentially expressed**. The figure displays the power and FDR from 100 simulations under homoscedastic scenario with two different sample sizes and 10% genes labeled as DE (pDE = 0.1). Panel (A) and (C) show the boxplot of powers from 7 DE methods arranged left to right in increasing values of median powers for $N = 40$ and $N = 100$, and, (B) and (D) show the FDR of the same simulation experiments. Methods corresponding to the boxplots on right side of the power plots are generally more efficient in finding DE genes. Methods that effectively manage to keep the FDR at or below the predetermined nominal level (0.05) should have their corresponding boxplot's median line positioned at or beneath the dotted red line.

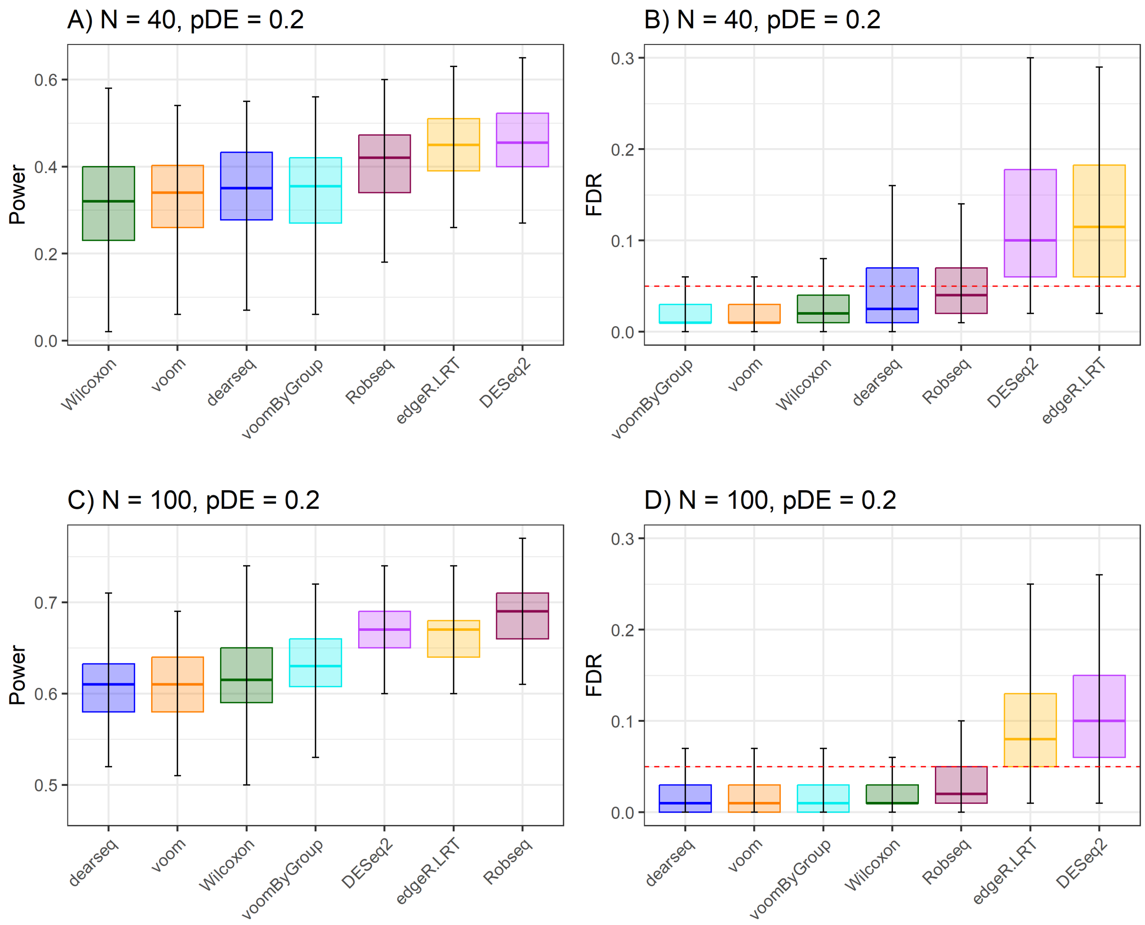

**Supplementary Figure** 6: **Power and FDR evaluations in heteroscedastic data when 20% genes were labeled as differentially expressed**. The figure displays the power and FDR from 100 simulations under heteroscedastic scenario with two different sample sizes and 20% genes labeled as DE (pDE = 0.2). Panel (A) and (C) show the boxplot of powers from 7 DE methods arranged left to right in increasing values of median powers for $N = 40$ and $N = 100$, and, (B) and (D) show the FDR of the same simulation experiments. Methods corresponding to the boxplots on right side of the power plots are generally more efficient in finding DE genes. Methods that effectively manage to keep the FDR at or below the predetermined nominal level (0.05) should have their corresponding boxplot's median line positioned at or beneath the dotted red line.

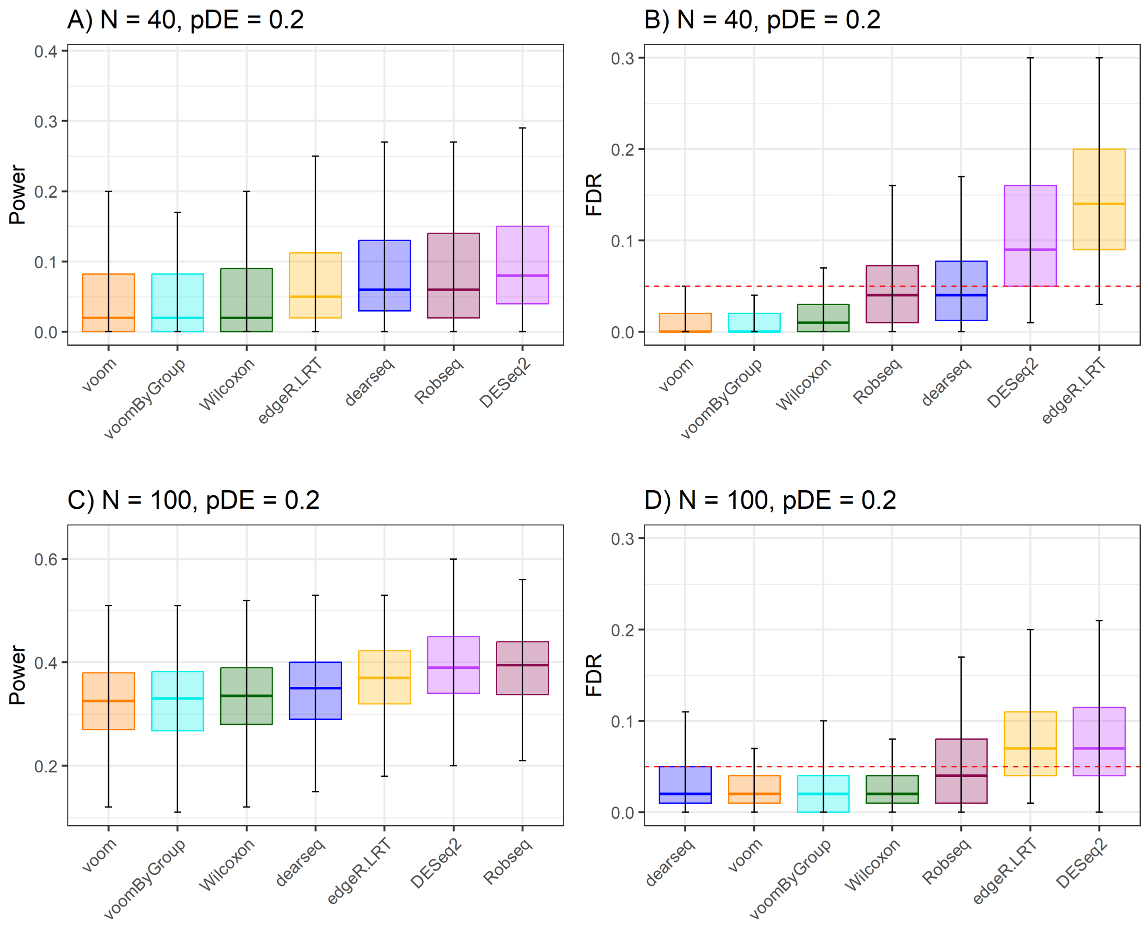

**Supplementary Figure 7: Power and FDR evaluations in homoscedastic data when 20% genes were labeled as differentially expressed.** The figure displays the power and FDR from 100 simulations under homoscedastic scenario with two different sample sizes and 20% genes labeled as DE (pDE = 0.2). Panel (A) and (C) show the boxplot of powers from 7 DE methods arranged left to right in increasing values of median powers for $N = 40$ and $N = 100$, and, (B) and (D) show the FDR of the same simulation experiments. Methods corresponding to the boxplots on right side of the power plots are generally more efficient in finding DE genes. Methods that effectively manage to keep the FDR at or below the predetermined nominal level (0.05) should have their corresponding boxplot's median line positioned at or beneath the dotted red line.

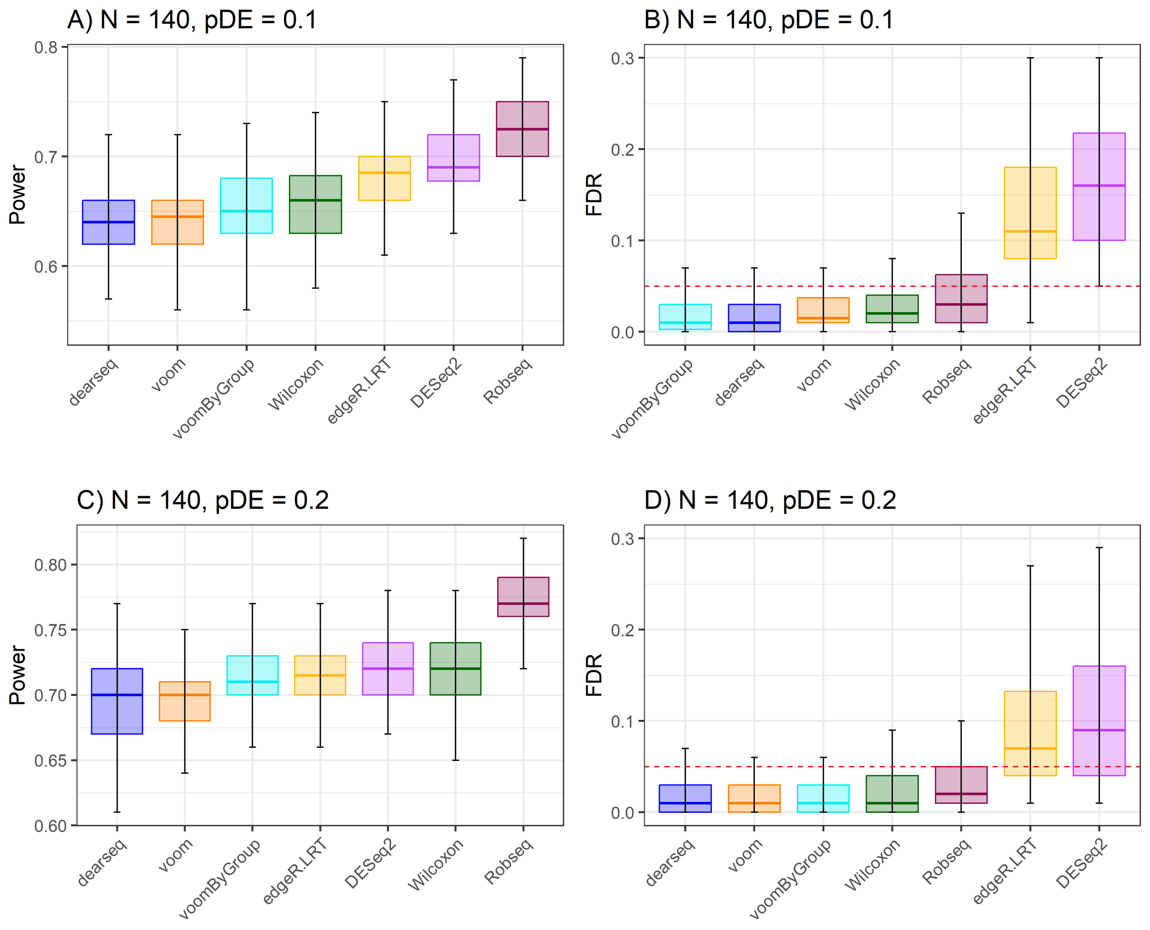

**Supplementary Figure** 8: **Power and FDR evaluations in heteroscedastic data when sample size N = 140.** The figure displays the power and FDR from 100 simulations under heteroscedastic scenario with 10% (pDE = 0.1) and 20% (pDE = 0.2) genes labeled as DE. Panel (A) and (C) show the boxplot of powers from 7 DE methods arranged left to right in increasing values of median powers for $N = 140$, and, (B) and (D) show the FDR of the same simulation experiments. Methods corresponding to the boxplots on right side of the power plots are generally more efficient in finding DE genes. Methods that effectively manage to keep the FDR at or below the predetermined nominal level (0.05) should have their corresponding boxplot's median line positioned at or beneath the dotted red line.

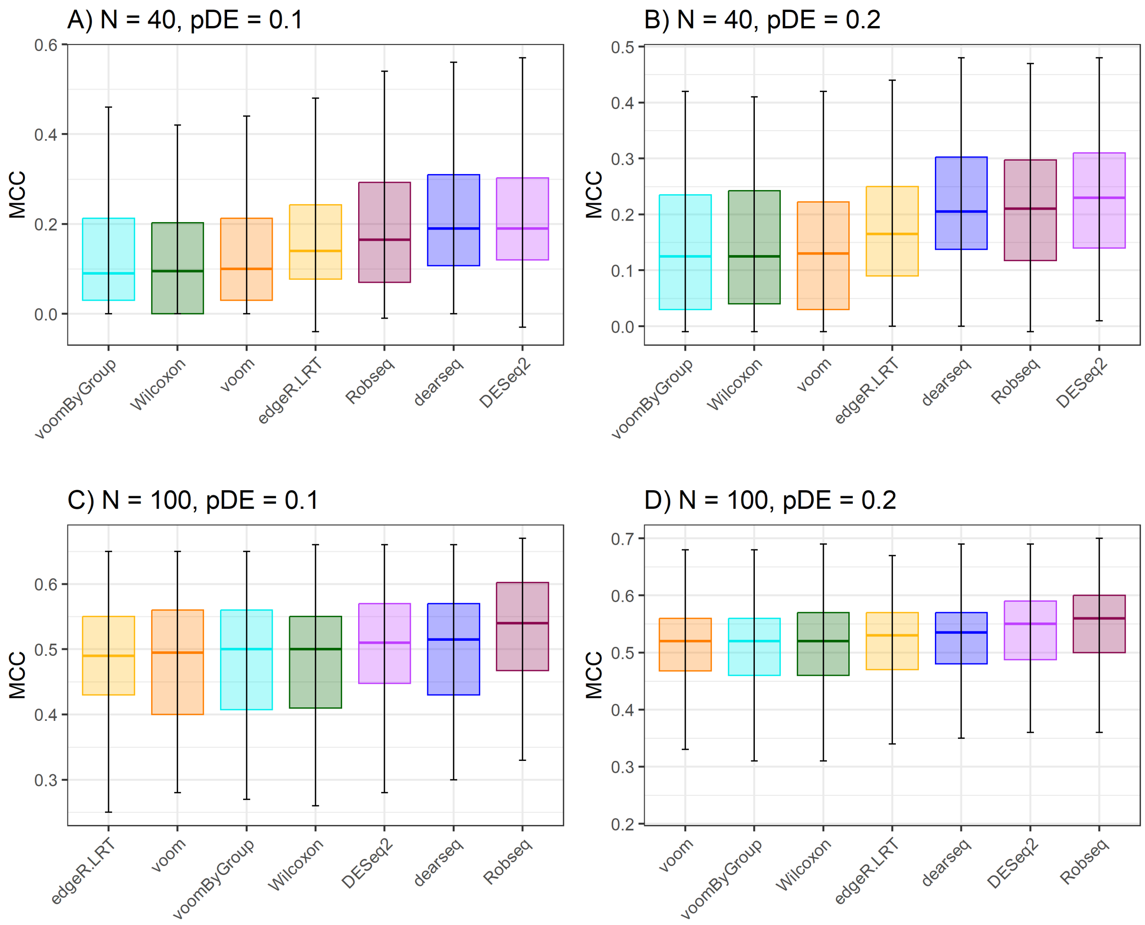

**Supplementary Figure** 9: **MCC score evaluations in homoscedastic data under varying sample size and proportion of DE genes**. The figure displays the MCC scores from 100 simulations under homoscedastic scenario with two sample sizes and two levels of proportions of DE genes (pDE). Panel (A) and (C) show the boxplot of MCC scores from 7 DE methods when the sample size N = 40 and 100 with true proportion of DE genes in the simulated dataset set at 10% (pDE = 0.1), and, (B) and (D) show the MCC scores of the same simulation experiments with the true proportion of DE genes set at 20% (pDE = 0.2). The boxplots are arranged in the increasing order of the median MCC scores from left to right in each plot. Methods with high MCC scores more accurately classify the genes in DE and non-DE categories.

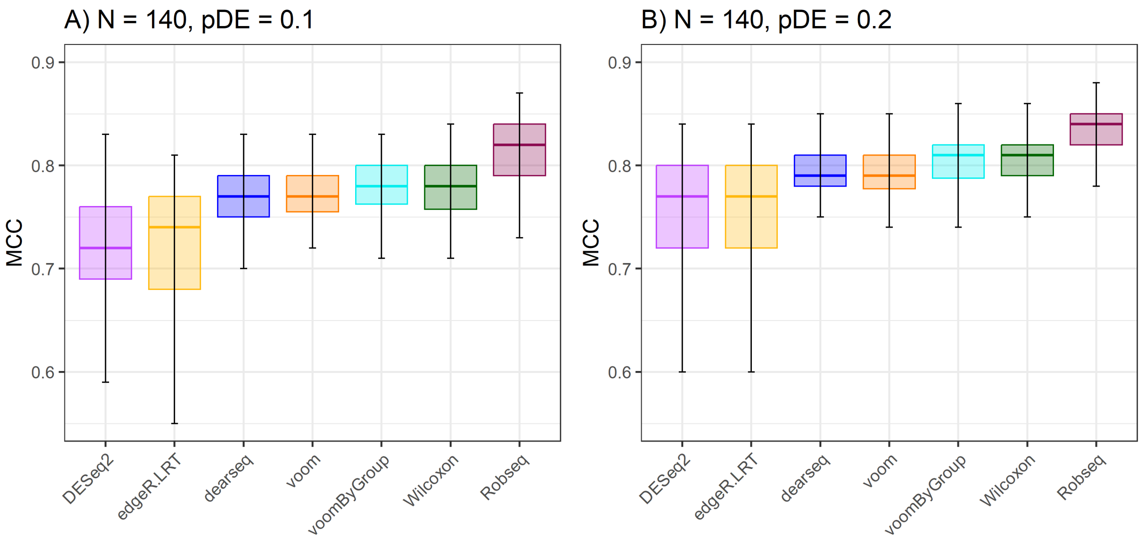

**Supplementary Figure** 10: **MCC score evaluations in heteroscedastic data under sample size N = 140 and varying proportion of DE genes**. The figure displays the MCC scores from 100 simulations under heteroscedastic scenario with sample size as $N = 140$ and two levels of proportions of DE genes (pDE). Panel (A) shows the boxplot of MCC scores from 7 DE methods when the true proportion of DE genes in the simulated dataset set at 10% (pDE = 0.1), and, (B) shows the MCC scores of the simulation experiments with the true proportion of DE genes set at 20% (pDE = 0.2). The boxplots are arranged in the increasing order of the median MCC scores from left to right in each plot. Methods with high MCC scores more accurately classify the genes in DE and non-DE categories.

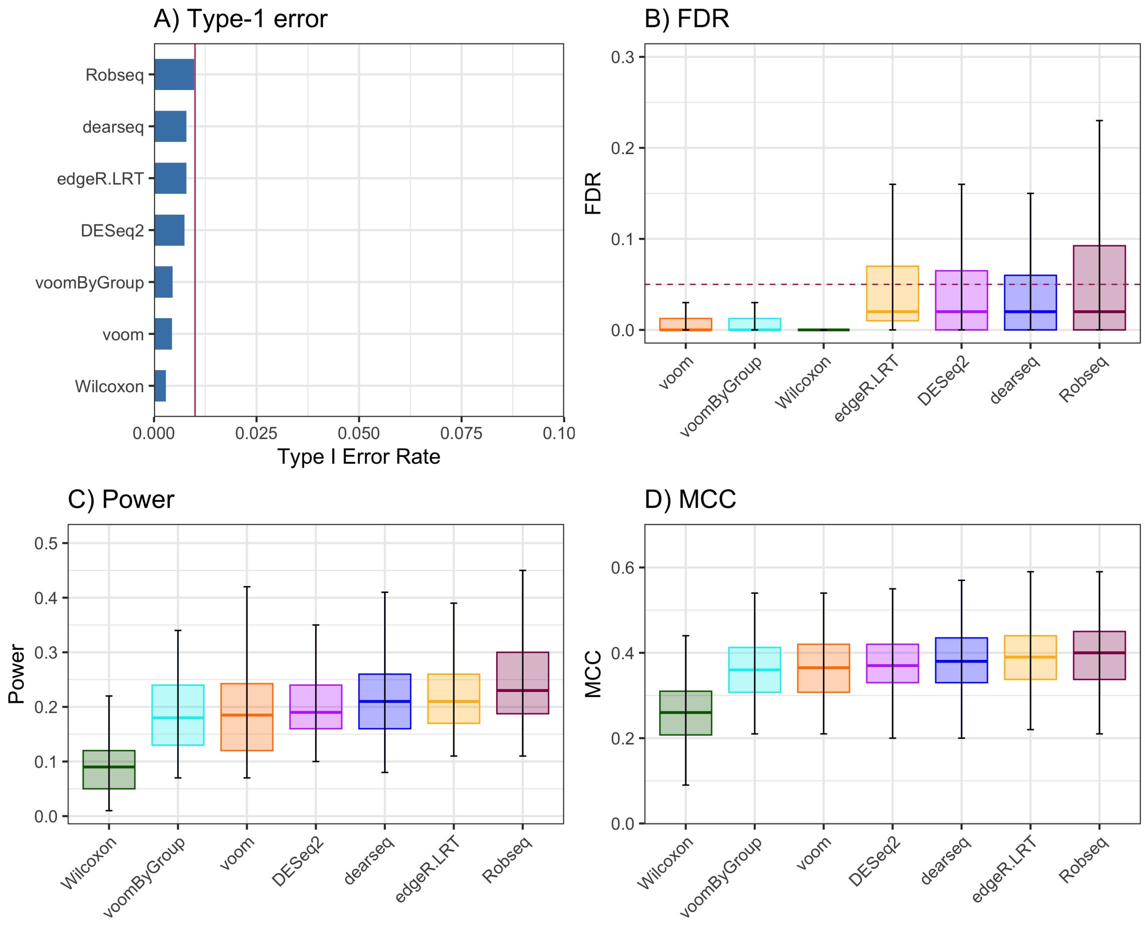

**Supplementary Figure** 11: **Evaluation of long non-coding RNA expression data under sample size N = 30 with 20% differentially expressed genes and type-1 error rate of 1%**. The figure shows the (A) type-I error, (B) FDR, (C) power, and (D) MCC scores from 100 simulation experiments based on a long non-coding RNA-Seq dataset using a total sample size of N = 30 and proportion of DE genes as 20% (pDE = 0.2). For panel (A), a red line in these plot represents the nominal type-1 error rate, set at $\alpha = 0.05$. These results are compiled from the median of 100 simulations. Methods that effectively maintain the type-1 error at or below this nominal threshold will have their corresponding bars positioned beneath the red line. Panel (B), shows the boxplot of FDR from 7 DE methods arranged left to right in increasing values of median FDRs. Methods that effectively manage to keep the FDR at or below the predetermined nominal level ($\alpha = 0.05$) should have their corresponding boxplot's median line positioned at or beneath the dotted red line. For panel (C), shows the boxplot of power from 7 DE methods arranged left to right in increasing values of median powers. Methods corresponding to the boxplots on right side of the power plot. For panel (D), boxplots are arranged in the increasing order of the median MCC scores from left to right in each plot. Methods with high MCC scores more accurately classify the genes in DE and non-DE categories.

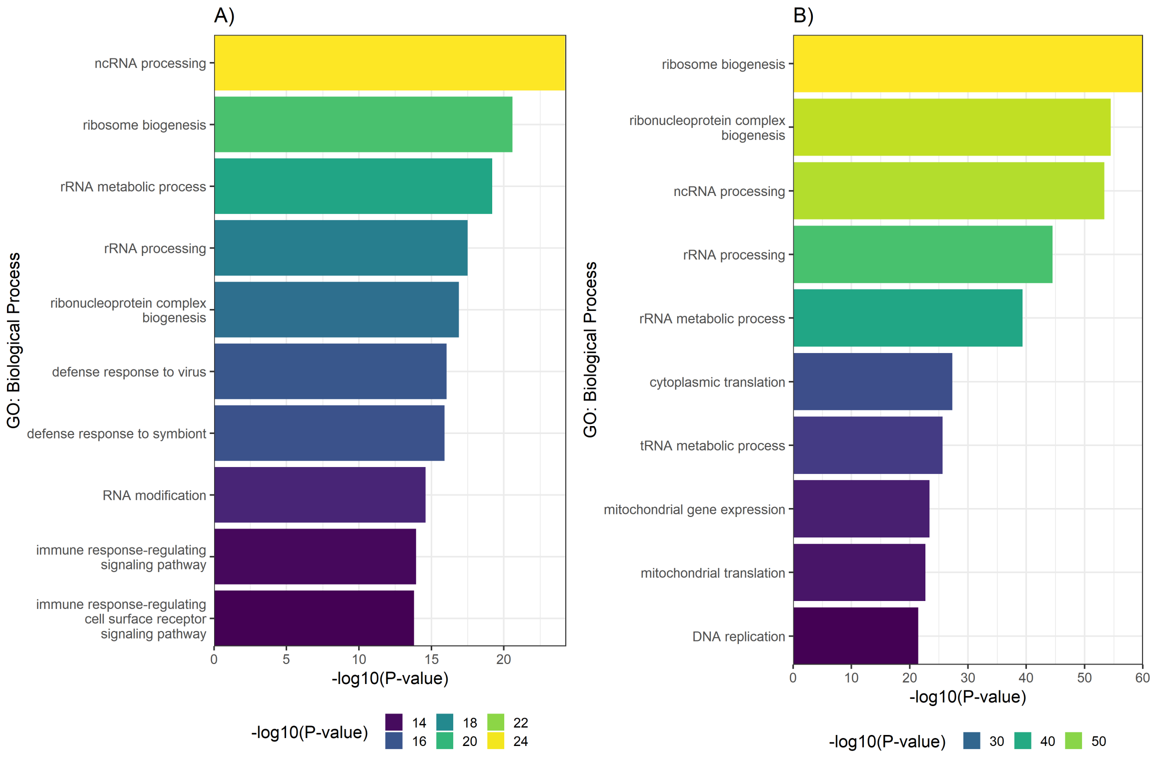

**Supplementary Figure** 12: **Robseq identified relevant pathways that were in consensus with other models**. In panel **(A)**, the bars represent the top ten enriched GO biological process terms associated with DE genes that were identified using Robseq and overlapped with other models in the lupus data. These top ten terms are selected based on having the lowest -log10 transformed p-values. The x-axis of the chart shows these -log10 transformed p-values for the various GO biological process terms. Additionally, a color scale is used in the chart, which varies according to the p-value of each term. In panel **(B)**, the bars represent the top ten enriched GO biological process terms associated with DE genes that were identified using Robseq and overlapped with other models in the colon cancer data. These top ten terms are selected based on having the lowest -log10 transformed p-values. The x-axis of the chart shows these -log10 transformed p-values for the various GO biological process terms. Additionally, a color scale is used in the chart, which varies according to the p-value of each term.

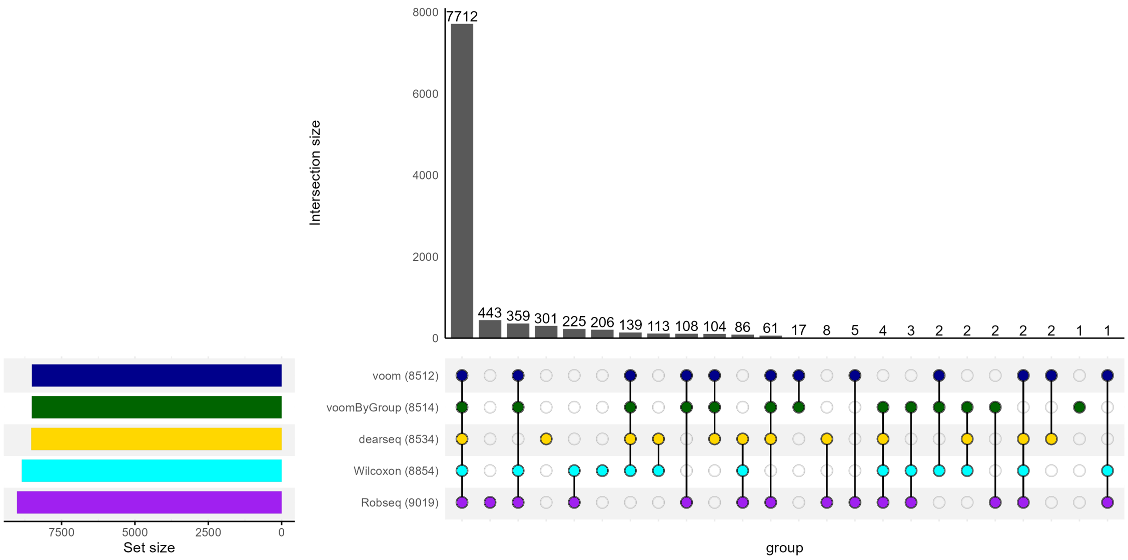

**Supplementary Figure** 13: **UpSet plot of number of DE genes detected across five DE models in the COPD data**. Using five DE models, we summarize the number of DE genes (out of 24,782 genes) detected between COPD (N = 98) and Normal (N = 91) groups in the COPD dataset. Numbers in parentheses represent the total number of DE genes identified by the corresponding method. Genes with observed FDR smaller than $\alpha = 0.05$ were deemed as DE.

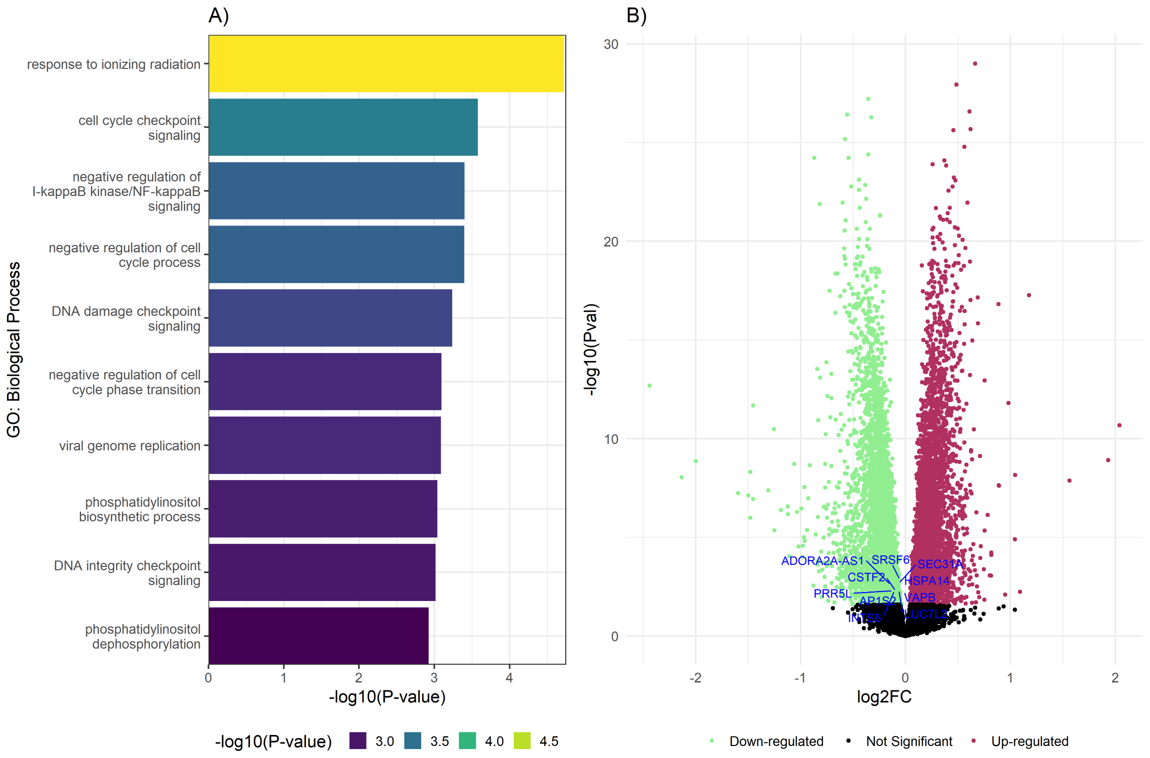

**Supplementary Figure** 14: **DNA damage and immunological pathways were found to be involved in COPD which were uniquely identified by Robseq**. In panel **(A)**, the bars represent the top ten enriched GO biological process terms associated with DE genes that were uniquely identified using Robseq. These top ten terms are selected based on having the lowest -log10 transformed p-values. The x-axis of the chart shows these -log10 transformed p-values for the various GO biological process terms. Additionally, a color scale is used in the chart, which varies according to the p-value of each term. In Panel **(B)**, a volcano plot illustrates the DE analysis of the COPD dataset conducted with Robseq. The x-axis of this plot displays the log2 fold change of genes analyzed in the COPD dataset, while the y-axis shows the -log10 transformed p-values of these genes. The points in green signify genes identified as DE with downregulation by Robseq, whereas the points in red indicate genes recognized as DE with upregulation by Robseq. The points in black represent genes that were not DE in the COPD dataset. The top ten DE genes, chosen for having the lowest FDR adjusted p-values and uniquely identified by Robseq, are highlighted with blue text.

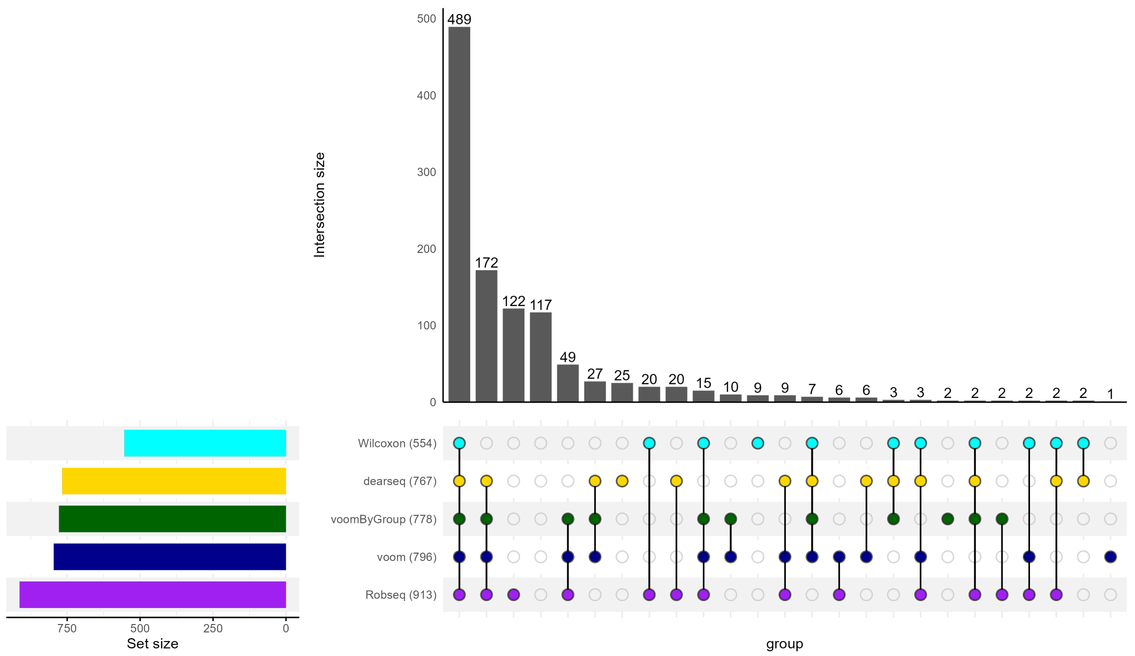

**Supplementary Figure** 15: **UpSet plot of number of DE lncRNAs detected across five DE models in the psoriasis data**. Using five DE models, we summarize the number of DE lncRNAs (out of 2959 lncRNAs) detected between healthy control (N = 16) and psoriasis patient (N = 36) groups in the psoriasis dataset. Numbers in parentheses represent the total number of DE lncRNAs identified by the corresponding method. LncRNAs with observed FDR smaller than $\alpha = 0.05$ were deemed as DE.

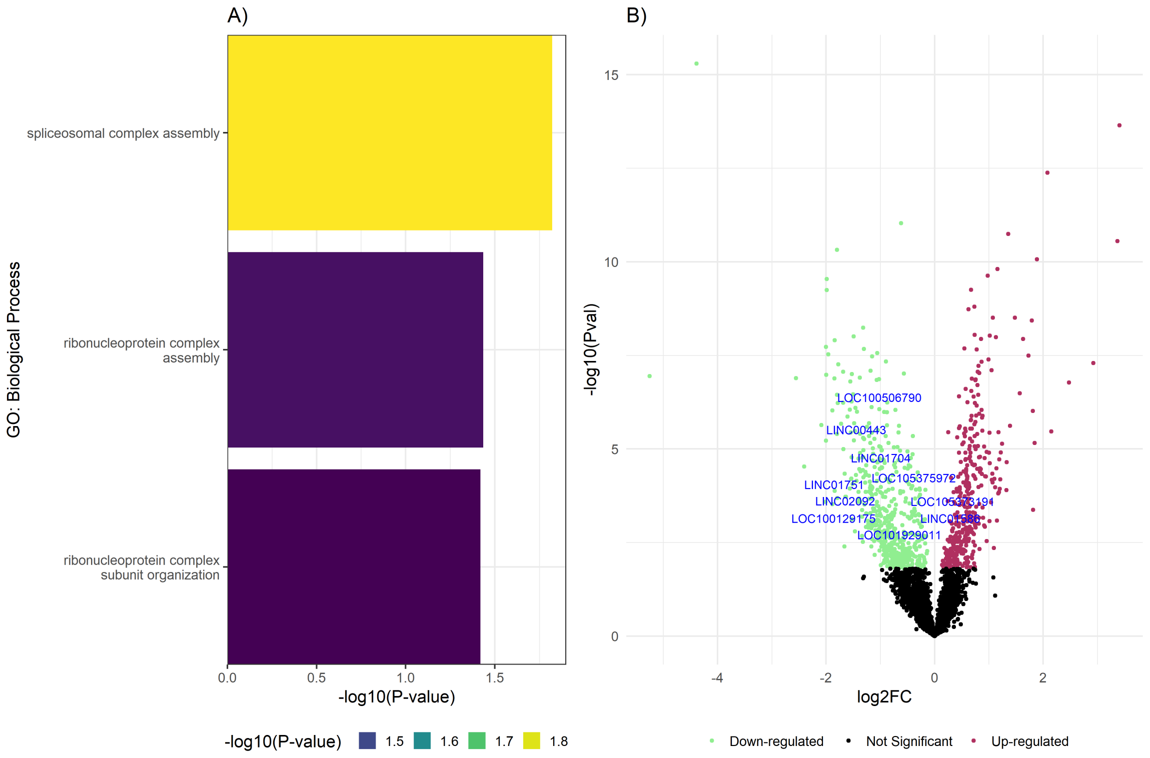

**Supplementary Figure** 16: **Spliceosomal complex and ribonucleoprotein complex assembly pathways were found to be involved in psoriasis which were uniquely identified by Robseq**. In panel **(A)**, the bars represent the enriched GO biological process terms associated with DE genes that were uniquely identified using Robseq. These terms were selected based on having the lowest -log10 transformed p-values. The x-axis of the chart shows these -log10 transformed p-values for the various GO biological process terms. Additionally, a color scale is used in the chart, which varies according to the p-value of each term. In Panel **(B)**, a volcano plot illustrates the DE analysis of the psoriasis dataset conducted with Robseq. The x-axis of this plot displays the log2 fold change of lncRNAs analyzed in the psoriasis dataset, while the y-axis shows the -log10 transformed p-values of these lncRNAs. The points in green signify lncRNAs identified as DE with downregulation by Robseq, whereas the points in red indicate lncRNAs recognized as DE with upregulation by Robseq. The points in black represent genes that were not found to be DE in the psoriasis dataset. The top ten DE lncRNAs, chosen for having the lowest FDR adjusted p-values and uniquely identified by Robseq, are highlighted with blue text.

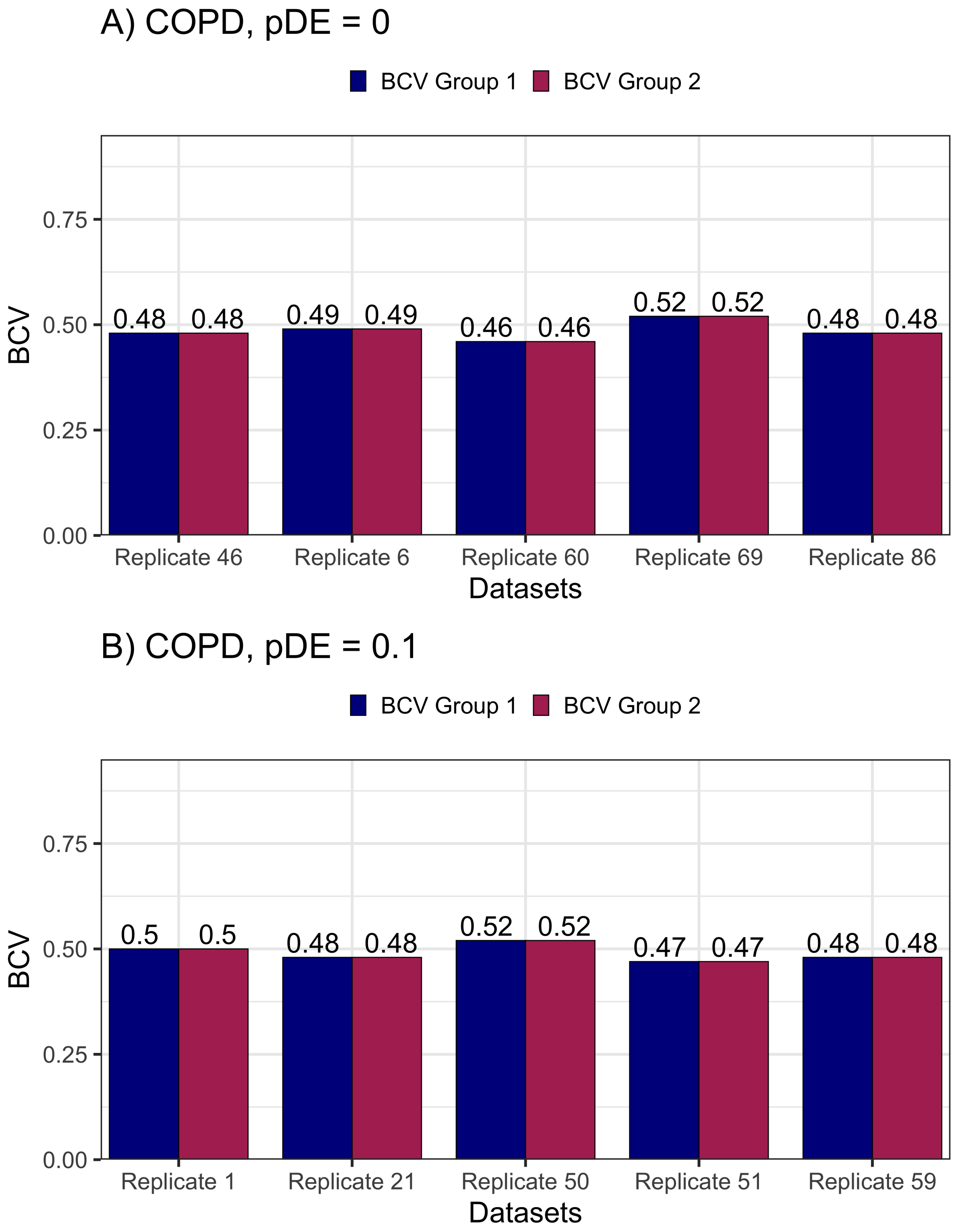

**Supplementary Figure** 17: **SimSeq accurately generates homoscedastic data when the input data features homoscedasticity.** This plot demonstrates that SimSeq produces homoscedastic synthetic data when provided with a homoscedastic template (COPD data). To confirm the homoscedastic nature of the synthetic data, we evaluated group-specific BCV values from five randomly selected replicates (out of 100 independent replicates) confirming their identicalness. In panel A), SimSeq was given the COPD data as input and tasked to create synthetic data without any DE genes (pDE = 0). The identicalness in group-specific BCV values across five randomly chosen replicates from this scenario indicated that SimSeq successfully generated homoscedastic data. In panel B) SimSeq used the same COPD data but was instructed to generate synthetic data containing 10% DE genes (pDE = 0.1). The identical group-specific BCV values from five randomly chosen replicates in this scenario also confirm SimSeq's capability to accurately produce homoscedastic data. Notably, the group-specific BCV values closely mimic those observed in the original COPD data (BCV 1 = BCV 2 = 0.5).

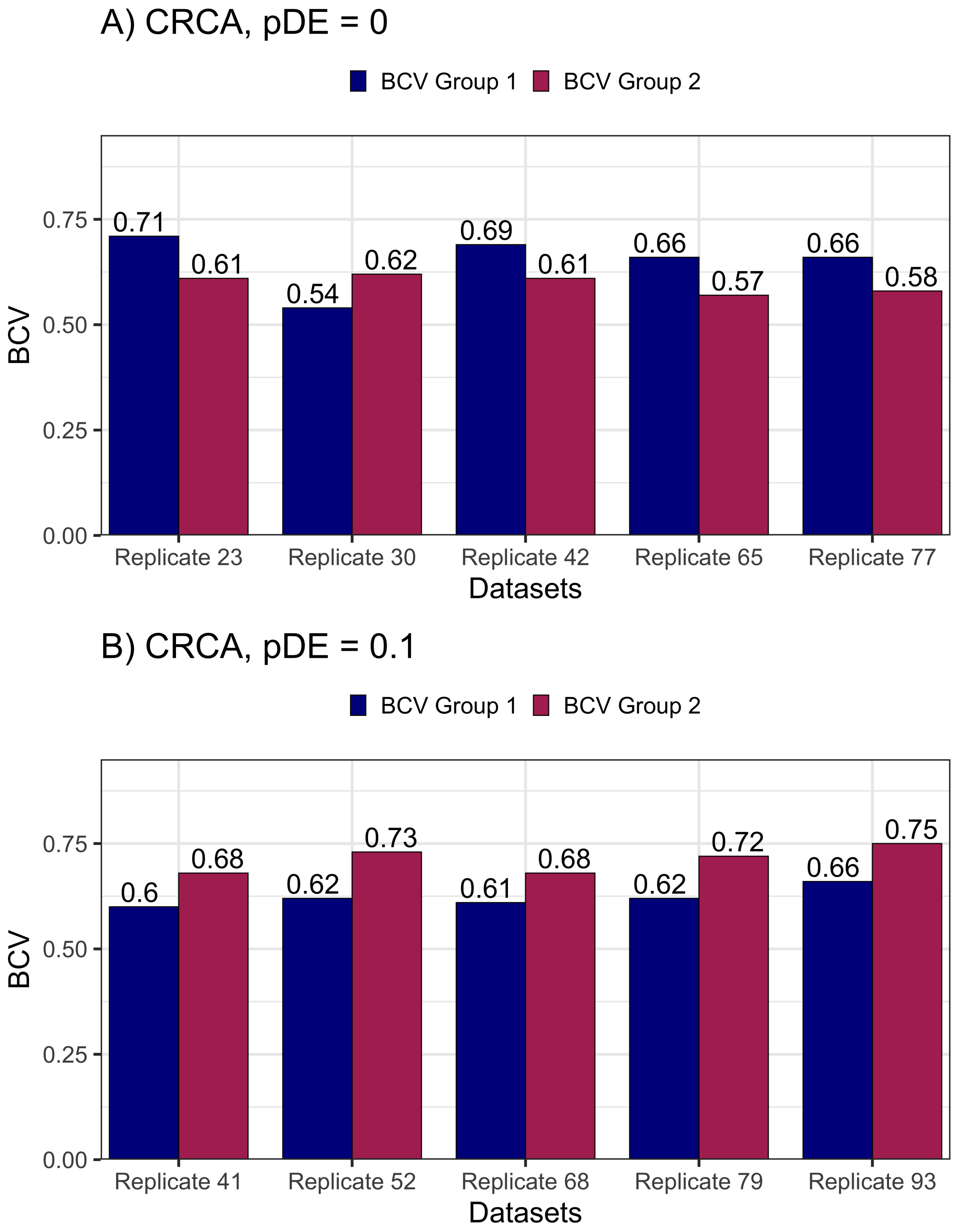

**Supplementary Figure** 18: **SimSeq accurately generates heteroscedastic data when the input data features heteroscedasticity.** This plot demonstrates that SimSeq produces heteroscedastic synthetic data when provided with a heteroscedastic template (CRCA data). To confirm the heteroscedastic nature of the synthetic data, we evaluated group-specific BCV values from five randomly selected replicates (out of 100 independent replicates) confirming their non-identicalness. In panel A),SimSeq was given the CRCA data as input and tasked to create synthetic data without any DE genes (pDE = 0). The variation in group-specific BCV values across five randomly chosen replicates from this scenario indicated that SimSeq successfully generated homoscedastic data. In panel B), SimSeq used the same CRCA data but was instructed to generate synthetic data containing 10% DE genes (pDE = 0.1). The variable group-specific BCV values from five randomly chosen replicates in this scenario also confirm SimSeq's capability to accurately produce homoscedastic data. Notably, the group-specific BCV values closely mimic those observed in the original CRCA data (BCV 1 = 0.66, BCV 2 = 0.73).

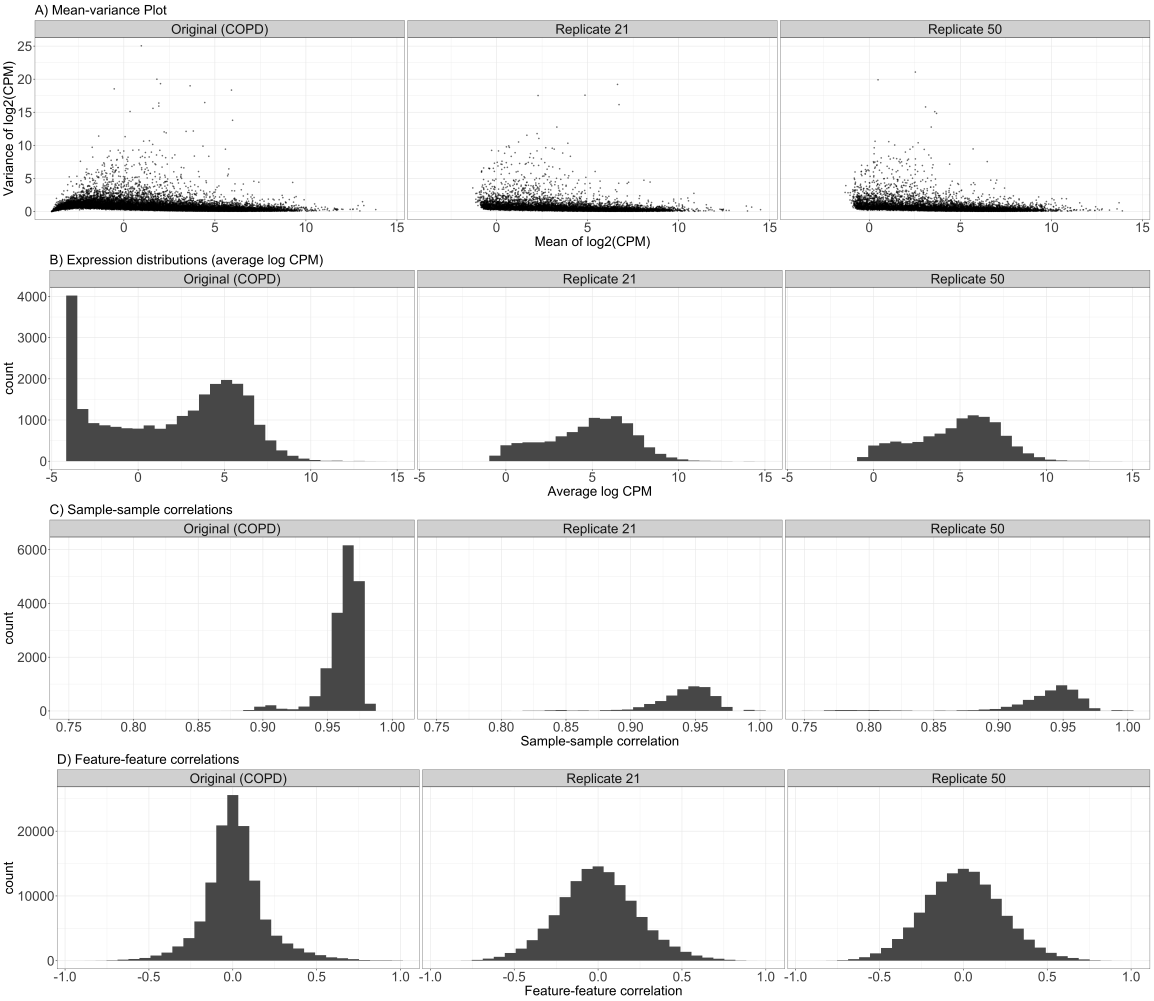

**Supplementary Figure** 19: **Quality assessment of homoscedastic data generated by Simseq.** This report compares two simulated count datasets generated using the SimSeq simulation procedure to the original COPD dataset. Each simulation comprises 100 independent runs with 10% true DE genes and 100 replicate sizes, with two randomly selected datasets presented for brevity. In A) These scatter plots depict the relationship between empirical mean and variance of features, based on log2(CPM) estimates across all samples. Unlike mean-dispersion plots, they disregard experimental design and sample grouping, solely focusing on overall mean and variance calculated using the cpm function from edgeR with a prior count of 2. B) These plots display the distribution of average abundance values for features, represented as log CPM values computed by edgeR. C) These plots display the distribution of Spearman correlation coefficients for sample pairs, derived from log(CPM) values using the cpm function from edgeR, with a prior count of 2. For datasets with over 500 samples, correlations are computed between 500 randomly selected pairs. D) These plots showcase the distribution of Spearman correlation coefficients between pairs of features, derived from log(CPM) values using the cpm function from edgeR with a prior count of 2. Only non-constant features are included, and if the dataset contains more than 500 such features, correlations between 500 randomly selected features are displayed. It's important to consider that the differences in characteristics could partially stem from the disparity in the dimensions of the synthetic data, which consists of 10,000 genes and 100 samples, compared to the original data, which includes 24,950 genes and 189 samples.

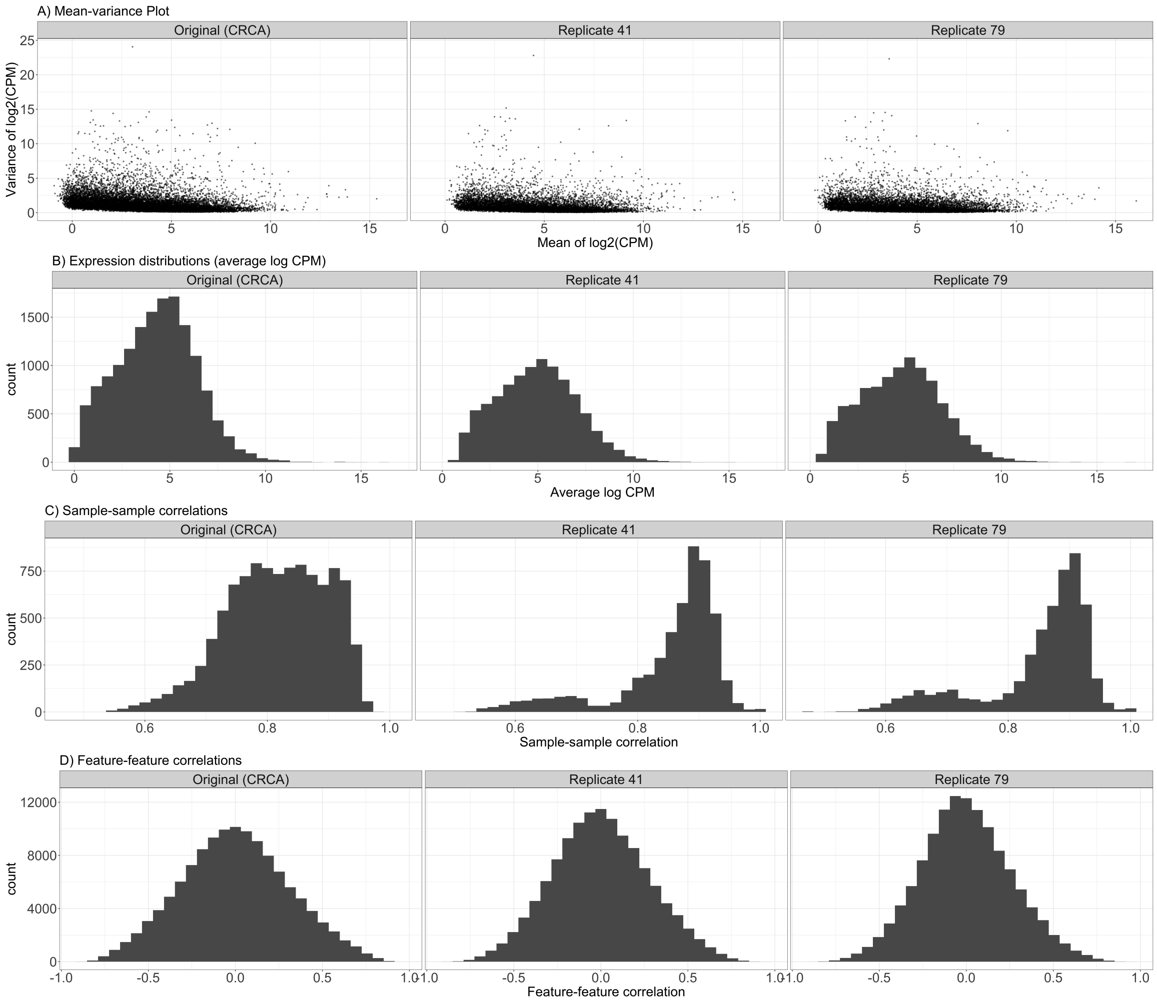

**Supplementary Figure** 20: **Quality assessment of heteroscedastic data generated by Simseq.** This report compares two simulated count datasets generated using the SimSeq simulation procedure to the original CRCA dataset. Each simulation comprises 100 independent runs with 10% true DE genes and 100 replicate sizes, with two randomly selected datasets presented for brevity. In A) These scatter plots depict the relationship between empirical mean and variance of features, based on log2(CPM) estimates across all samples. Unlike mean-dispersion plots, they disregard experimental design and sample grouping, solely focusing on overall mean and variance calculated using the cpm function from edgeR with a prior count of 2. B) These plots display the distribution of average abundance values for features, represented as log CPM values computed by edgeR. C) These plots display the distribution of Spearman correlation coefficients for sample pairs, derived from log(CPM) values using the cpm function from edgeR, with a prior count of 2. For datasets with over 500 samples, correlations are computed between 500 randomly selected pairs. D) These plots showcase the distribution of Spearman correlation coefficients between pairs of features, derived from log(CPM) values using the cpm function from edgeR with a prior count of 2. Only non-constant features are included, and if the dataset contains more than 500 such features, correlations between 500 randomly selected features are displayed. It's important to consider that the differences in characteristics could partially stem from the disparity in the dimensions of the synthetic data, which consists of 10,000 genes and 100 samples, compared to the original data, which includes 27,284 genes and 144 samples.

#### 2 Supplementary Tables

**Supplementary Table** 1: This table presents 12 bulk RNA-Seq datasets across various human diseases and tissues, as utilized in Figure 1 to investigate group heteroscedasticity. Each row provides specific details regarding the datasets. The first column indicates the source from which the data was obtained. The second column lists the dataset names. The third and fourth columns denote the condition labels for the first and second experimental/disease groups, respectively. The fifth column displays the number of samples within each dataset, while the sixth column indicates the number of genes included in the analysis. The seventh column contains the Pubmed ID associated with each dataset, and the final column provides the Accession no. or the web links to access these data.

| Data Source | Data Name | Condition 1 | Condition 2 | Number of Samples | Number of Genes | Pubmed | Accession |
|---|---|---|---|---|---|---|---|
| **GEO** | Polycystic Ovary | Control | Polycystic ovary | 12 | 19248 | 34552933 | GSE155489 |
| | Heart Failiure | Heart failure | Healthy donor | 30 | 26289 | 31902369 | GSE135055 |
| | Bipolar | Neurotypical | Bipolar disorder | 33 | 24572 | 33667413 | GSE157509 |
| | Obesity | Normal-weight child with asthma | Obese child with asthma | 42 | 30277 | 24733242 | GSE86430 |
| | Cirrhosis | Healthy human liver | Cirrhotic NASH* human liver | 45 | 24304 | 33901295 | GSE180882 |
| | Alzheimer | Alzheimer's disease | Control | 117 | 24281 | 29346778 | GSE95587 |
| | Lupus | Systemic lupus erythematosus | Healthy | 120 | 24408 | 31890206 | GSE112087 |
| | Colon Cancer | Native tissue | Tumor tissue | 144 | 27248 | 34737287 | GSE156451 |
| **TCGA** | Liver Cancer | Normal tissue | Tumor tissue | 100 | 52196 | | GDC Xena Hub |
| | Lung Cancer | Normal tissue | Tumor tissue | 114 | 54140 | | |
| **GTEx** | Prostate & Brain | Prostate | Brain cortex | 426 | 54235 | | GTEx Portal |
| | Heart | Heart - atrial appendage | Heart - left ventricle | 758 | 53950 | | |

* NASH: nonalcoholic steatohepatitis

**Supplementary Table** 2: This table displays the top ten genes with the lowest FDR adjusted p-values (top hits), which were uniquely identified by Robseq, in the analysis of Lupus data. The first column lists the gene symbols, the second column displays the estimated $Log_2$ fold change for Robseq for each respective gene, the third column shows the standard error, the fourth column indicates the lower confidence interval, the fifth column presents the upper confidence interval, the sixth column denotes the significance level, and the last column represents the multiple testing adjusted p-value.

| **Genes** | $Log_2$ **Fold Change** | **Standard Error** | **Lower Confidence Interval** | **Upper Confidence Interval** | **p-value** | **Adjusted p-value** |
|---|---|---|---|---|---|---|
| SP140L | 0.113 | 0.024 | 0.160039136 | 0.065960864 | 5.73E-06 | 3.12E-05 |
| BTNL3 | -1.838 | 0.416 | -1.022654982 | -2.653345018 | 2.36E-05 | 0.000107646 |
| LINC01138 | 0.094 | 0.023 | 0.139079172 | 0.048920828 | 8.80E-05 | 0.000341504 |
| EIF2AK1 | -0.309 | 0.08 | -0.152202881 | -0.465797119 | 0.000177317 | 0.000635841 |
| FCN1 | 0.179 | 0.047 | 0.271118307 | 0.086881693 | 0.000263962 | 0.000898711 |
| TBC1D7 | 0.11 | 0.03 | 0.16879892 | 0.05120108 | 0.000337507 | 0.001118614 |
| FECH | -0.424 | 0.124 | -0.180964466 | -0.667035534 | 0.000843461 | 0.002503815 |
| MPP1 | -0.335 | 0.098 | -0.14292353 | -0.52707647 | 0.000846514 | 0.002511422 |
| MYO5C | 0.185 | 0.055 | 0.292798019 | 0.077201981 | 0.000967648 | 0.002816409 |
| ARID1A | 0.143 | 0.043 | 0.227278451 | 0.058721549 | 0.001119879 | 0.003191758 |

**Supplementary Table** 3: This table displays the top ten genes with the lowest FDR adjusted p-values (top hits), which were uniquely identified by Robseq, in the analysis of colon Cancer data. The first column lists the gene symbols, the second column displays the estimated $Log_2$ fold change for Robseq for each respective gene, the third column shows the standard error, the fourth column indicates the lower confidence interval, the fifth column presents the upper confidence interval, the sixth column denotes the significance level, and the last column represents the multiple testing adjusted p-value.

| **Genes** | $Log_2$ **Fold Change** | **Standard Error** | **Lower Confidence Interval** | **Upper Confidence Interval** | **p-value** | **Adjusted p-value** |
|---|---|---|---|---|---|---|
| CDC42BPB | -0.156 | 0.038 | -0.081521369 | -0.230478631 | 7.50E-05 | 0.00013789 |
| SLC10A5 | -0.444 | 0.112 | -0.224484034 | -0.663515966 | 0.000119271 | 0.000214677 |
| EXOC8 | -0.182 | 0.047 | -0.089881693 | -0.274118307 | 0.000146151 | 0.000260901 |
| ZBTB34 | -0.207 | 0.054 | -0.101161945 | -0.312838055 | 0.000183504 | 0.000323417 |
| COX1 | -0.517 | 0.139 | -0.244565006 | -0.789434994 | 0.000300124 | 0.000517178 |
| SMPD2 | -0.202 | 0.055 | -0.094201981 | -0.309798019 | 0.000301302 | 0.000519125 |
| TMEM117 | -0.293 | 0.08 | -0.136202881 | -0.449797119 | 0.000339322 | 0.000581113 |
| PPP2R5E | -0.242 | 0.067 | -0.110682413 | -0.373317587 | 0.000400133 | 0.000679607 |
| LRP4-AS1 | -0.725 | 0.201 | -0.331047239 | -1.118952761 | 0.00043824 | 0.000740781 |
| ICMT | -0.174 | 0.048 | -0.079921729 | -0.268078271 | 0.000443378 | 0.000749135 |

**Supplementary Table** 4: This table displays the top ten genes with the lowest FDR adjusted p-values (top hits), which were uniquely identified by Robseq, in the analysis of COPD data. The first column lists the gene symbols, the second column displays the estimated $Log_2$ fold change for Robseq for each respective gene, the third column shows the standard error, the fourth column indicates the lower confidence interval, the fifth column presents the upper confidence interval, the sixth column denotes the significance level, and the last column represents the multiple testing adjusted p-value.

| **Genes** | $Log_2$ **Fold Change** | **Standard Error** | **Lower Confidence Interval** | **Upper Confidence Interval** | **p-value** | **Adjusted p-value** |
|---|---|---|---|---|---|---|
| TUBGCP2 | -0.087 | 0.022 | -0.043880792 | -0.130119208 | 9.50E-05 | 0.000352762 |
| DKC1 | -0.099 | 0.026 | -0.048040936 | -0.149959064 | 0.00024957 | 0.00084506 |
| HNRNPL | -0.056 | 0.016 | -0.024640576 | -0.087359424 | 0.000758058 | 0.002269199 |
| YPEL5 | -0.096 | 0.028 | -0.041121008 | -0.150878992 | 0.000866918 | 0.002559316 |
| ZNF527 | -0.117 | 0.035 | -0.048401261 | -0.185598739 | 0.00088021 | 0.00259425 |
| RIMKLB | -0.113 | 0.034 | -0.046361225 | -0.179638775 | 0.000930553 | 0.00272816 |
| C5orf42 | -0.121 | 0.036 | -0.050441297 | -0.191558703 | 0.001084761 | 0.003125157 |
| DIEXF | -0.084 | 0.025 | -0.0350009 | -0.1329991 | 0.001184102 | 0.00338285 |
| SRSF6 | -0.059 | 0.018 | -0.023720648 | -0.094279352 | 0.001246794 | 0.003546006 |
| TRIO | -0.09 | 0.028 | -0.035121008 | -0.144878992 | 0.001260937 | 0.003581078 |

**Supplementary Table** 5: This table displays the top ten genes with the lowest FDR adjusted p-values (top hits), which were uniquely identified by Robseq, in the analysis of psoriasis data. The first column lists the gene symbols, the second column displays the estimated $Log_2$ fold change for Robseq for each respective gene, the third column shows the standard error, the fourth column indicates the lower confidence interval, the fifth column presents the upper confidence interval, the sixth column denotes the significance level, and the last column represents the multiple testing adjusted p-value.

| **Genes** | $Log_2$ **Fold Change** | **Standard Error** | **Lower Confidence Interval** | **Upper Confidence Interval** | **p-value** | **Adjusted p-value** |
|---|---|---|---|---|---|---|
| LOC100506790 | -1.16 | 0.182 | -0.803286555 | -1.516713445 | 7.63E-07 | 2.97E-05 |
| LINC00443 | -1.3 | 0.227 | -0.855088176 | -1.744911824 | 5.52E-06 | 0.00012192 |
| LINC01704 | -1.137 | 0.205 | -0.735207383 | -1.538792617 | 9.70E-06 | 0.000188853 |
| LOC105375972 | -0.989 | 0.216 | -0.565647779 | -1.412352221 | 0.000104608 | 0.001097641 |
| LINC01751 | -1.206 | 0.275 | -0.667009904 | -1.744990096 | 0.000137568 | 0.001325318 |
| LOC105373191 | -0.529 | 0.124 | -0.285964466 | -0.772035534 | 0.000209363 | 0.001838291 |
| LINC02092 | -1.007 | 0.252 | -0.513089076 | -1.500910924 | 0.000403132 | 0.003155734 |
| LINC01586 | -0.352 | 0.092 | -0.171683313 | -0.532316687 | 0.00060268 | 0.004293998 |
| LOC100129175 | -0.912 | 0.247 | -0.427888896 | -1.396111104 | 0.000895415 | 0.005772406 |
| LOC101929011 | -0.786 | 0.217 | -0.360687815 | -1.211312185 | 0.001096461 | 0.00674517 |

### 3 Supplementary Text

#### 3.1 Population-level analysis of COPD dataset with Robseq

The first additional dataset we examined in our analysis was a COPD dataset, consisting of 24,782 gene expression profiles across 189 samples. Chronic Obstructive Pulmonary Disease (COPD) is a severe condition characterized by airflow limitation, leading to breathing difficulties and impaired lung function [9]. Research indicates COPD is genetically driven, involving alterations in various genomic regions identified through genome-wide association studies (GWAS) and Mendelian syndromes such as alpha-1 antitrypsin deficiency [10]. Several genes and proteins related to COPD have been identified, playing key roles in disease susceptibility, progression, and heterogeneity, including those involved in lung parenchymal destruction (emphysema), airway disease, and clinical manifestations such as exacerbation frequency and exercise capacity. However, the precise molecular mechanisms of COPD remain elusive, necessitating further research [9]. Among other features, this dataset showed group homoscedasticity, evidenced by an identical BCV of 0.5 in both the healthy and the COPD group. Previous analyses of this dataset did not account for the inherent group homoscedasticity, so we re-analyzed it using a model like Robseq, which considers group homoscedasticity, to potentially gain better biological insights.

In our analysis of COPD data, Robseq identified a total of 9019 DE genes, which was the highest number compared to all other models. Additionally, there was a significant overlap of these genes (7712 in total) with those identified by other models (**Fig. S13**). Notably, Robseq uniquely detected 443 DE genes that were not identified by other models. We have listed the top ten genes with the lowest FDR adjusted p-values (top hits), which were exclusively identified by Robseq, in **Table S4** and in **Fig. S14B**. The GO analysis showed that the DE genes overlapping with other models were primarily involved in ribonucleoprotein complex biogenesis, ribosome biogenesis, mitochondrial translation, and RNA splicing. In contrast, the DE genes uniquely identified by Robseq were associated with pathways related to DNA damage, cell cycle checkpoint, nucleoplasm, immunological pathways, and innate immunity (**Fig S14A**). Robseq uniquely identified certain genes, notably emphasizing immune-related pathways. This included the identification of DE genes like CCR4, a receptor found on innate immune cells for CCL17/CCL22 [13], and FGL2, a protein expressed by macrophages, T cells, and tumor cells with roles in coagulation and immune suppression [12,4]. Additionally, Robseq identified 16 unique DE antisense RNAs, although their functional connections to diseases are largely unknown. For example, WDFY3-AS2 and ADORA2A-AS1 are linked to cancer and immunological pathways, suggesting that other identified RNAs may have new roles in COPD [5,2]. These findings collectively underscore Robseq's effectiveness in unveiling novel and relevant biological mechanisms in both heteroscedastic and homoscedastic scenarios.

#### 3.2 Population-level analysis of psoriasis dataset with Robseq

In this additional data analysis a psoriasis long non-coding RNA-Seq dataset was used, consisting of 2959 lncRNA expression profiles across 52 samples. Psoriasis is a chronic inflammatory skin disease characterized by varying degrees of severity, ranging from mild to severe, often causing significant discomfort and distress to affected individuals [1]. Research indicates psoriasis is genetically driven, involving alterations in DNA segments such as single nucleotide polymorphisms (SNPs), copy number variants (CNVs), and copy-neutral loss of heterozygosity (LOH), contributing to disease susceptibility [11]. Several genes and proteins related to psoriasis have been identified, playing key roles in immune dysregulation, epidermal cell proliferation, and differentiation [1]. These include genes involved in the interleukin signaling pathway, epidermal growth factor receptor (EGFR) signaling, and keratinocyte differentiation. However, the precise molecular mechanisms of psoriasis remain elusive, with ongoing research aimed at unraveling the underlying disease pathology [1]. Among other features, this dataset showed group heteroscedasticity, evidenced by a BCV of 0.4 in the healthy group compared to 0.56 in the SLE group. Previous analyses of this dataset did not account for the inherent group heteroscedasticity, so we re-analyzed it using a model like Robseq, which considers group heteroscedasticity, to potentially gain better biological insights.

In our analysis of psoriasis long non-coding RNA-Seq data, Robseq identified a total of 913 DE genes, which was the highest number compared to all other models. Additionally, there was a significant overlap of these genes (489 in total) with those identified by other models (**Fig. S15**). Notably,

Robseq uniquely detected 122 DE genes that were not identified by other models. We have listed the top ten genes with the lowest FDR-adjusted p-values (top hits), which were exclusively identified by Robseq (**Table S5 - Fig. S16B**). The gene ontology (GO) analysis showed that the DE genes overlapping with other models were primarily involved in the negative regulation of SMAD protein signal transduction, positive regulation of vascular-associated smooth muscle cell migration, and negative regulation of angiogenesis. In contrast, the DE genes uniquely identified by Robseq were associated with pathways related to the spliceosomal complex assembly, ribonucleoprotein complex assembly, and ribonucleoprotein complex subunit organization (**Fig. S16A**).

Most of the non-coding DE signals identified by Robseq are novel lncRNAs with unexplored functions. Among these, LINC00443 and OVAAL, which are associated with cancer, were found to be downregulated in psoriasis datasets. LINC00443, known for its tumor-suppressing function through competing endogenous RNA mechanisms, is downregulated in clear cell renal cell carcinoma [6,15]. However, the specific impact of its downregulation in psoriasis remains unclear due to a lack of molecular evidence. Conversely, OVAAL, which is also downregulated, plays a dual role in controlling RAF/MEK/ERK signaling and p27-mediated cell senescence, aiding in cancer cell survival [8]. This signaling pathway is crucial in the epidermis, with RAF1 playing a significant role in the development and progression of RAS-induced tumors by inhibiting keratinocyte differentiation, and MEK/ERK being involved in epidermal proliferation [7]. The reduced levels of OVAAL in psoriasis suggest a possible compromise in the RAF/MEK/ERK pathway, which may be linked to the development of the disease. In contrast, lncRNAs such as PCAT18 and LGALS8-AS1, typically upregulated in cancers, are also found to be elevated in psoriasis. PCAT18 is essential for the invasion, migration, and proliferation of castration-resistant prostate cancer cells [3]. LncRNA LGALS8-AS1, highly expressed in breast cancer, upregulates SOX12 by sponging miR-125b-5p and activating the PI3K/AKT signaling pathway [14]. The discovery of these lncRNAs in psoriasis by Robseq demonstrates its effectiveness not only in identifying mRNAs but also in uncovering biologically relevant lncRNAs.
